# Alternative exon splicing reveals hidden mitochondrial targeting of PLIN3

**DOI:** 10.64898/2026.02.18.706587

**Authors:** Axel KF Aguettaz, Yoan Arribat, Alexandra Schmitt, Aminata Gerihanov, Jocelyn Fleurimont, Jean Daraspe, Vithusan Vijayatheva, Cassandra Tabasso, Sylviane Lagarrigue, Ernesto Picardi, Sarah Cohen, Christel Genoud, Francesca Amati

## Abstract

Perilipin 3 (PLIN3) is a lipid droplet–associated protein implicated in cellular lipid metabolism. We characterize a previously unrecognized alternative splicing event in human *PLIN3*. The generated transcript *PLIN3B* is detectable at the RNA level across human tissues and is modulated by antisense oligonucleotide–mediated splicing. Despite clear evidence for *PLIN3B* transcript expression, endogenous PLIN3B protein could not be detected using multiple complementary state-of-the-art detection strategies.

When expressed exogenously, PLIN3B coding sequence encodes an unstable protein that localizes predominantly within mitochondria, in contrast to canonical PLIN3. Using super-resolution microscopy, correlative light–electron microscopy, and interaction proteomics, we show that PLIN3B accumulates intramitochondrially and engages protein quality control machinery. Mitochondrial targeting depends on residues shared by both isoforms, not altered by alternative splicing, indicating that mitochondrial targeting is normally repressed.

Together, these results show that alternative exon splicing in PLIN3 reveals latent mitochondrial targeting without producing detectable endogenous protein, underscoring the need to distinguish transcript diversity from stable protein expression.

## Introduction

Lipid droplets (LD) are dynamic organelles involved in lipid synthesis, breakdown, and trafficking, integrating lipid storage with cellular energy metabolism and organelle communication. LD formation, turnover, and interactions with the endoplasmic reticulum (ER) and mitochondria are tightly regulated by specialized protein networks that adapt LD behavior to metabolic state (Mathiowetz and Olzmann, 2024; Olzmann and Carvalho, 2019). Members of the perilipin (PLIN) protein family play central roles in these processes by controlling LD surface properties and mediating organelle interaction.

Perilipin 3 (PLIN3, also known as TIP47) is a ubiquitously expressed PLIN. Like most PLINs (Bickel et al., 2009), PLIN3 contains an N-terminal PAT domain, a central 11 mer repeats that form a membrane binding amphipathic helix (Rowe et al., 2016), and a C-terminal four-helix bundle (4HB) domain, which regulates LD binding (Ajjaji et al., 2019). PLIN3 has been shown to undergo regulated degradation through chaperone-mediated autophagy (CMA) (Kaushik and Cuervo, 2015), linking its turnover to cellular quality-control pathways.

Alternative splicing is a major source of functional diversity in mammalian proteomes and is increasingly recognized as a mechanism for modulating protein localization, stability, and function, including for members of the PLIN protein family (Hsieh et al., 2012). While investigating PLIN3 biology, we identified an alternative PLIN3 transcript generated by a previously undescribed splicing event affecting exon 6. This splicing event produces a shortened coding sequence, here referred to as PLIN3B. Although the corresponding exon is present in annotated transcripts, it has been classified as part of an incomplete coding sequence, and the existence of a stable endogenous protein product has not been established.

In this study, we systematically characterize the PLIN3B splice variant using molecular cloning, imaging, biochemical fractionation, proteomics, antisense-mediated splicing modulation, and analysis of public datasets. We show that the PLIN3B coding sequence, when expressed exogenously, localizes at mitochondria, in contrast to the LD- and cytosol-associated localization of PLIN3A. Importantly, despite clear evidence for *PLIN3B* transcript expression and efficient experimental induction of the splicing event, we find no evidence for endogenous PLIN3B protein expression. Together, our findings highlight the need for rigorous experimental discrimination between transcript diversity and stable protein expression and illustrate how alternative splicing can generate coding sequences that alter intracellular targeting without yielding detectable endogenous protein products.

## Results

### Identification of PLIN3B, a novel PLIN3 splicing isoform

While cloning *PLIN3* cDNA from human muscle samples, we identified two transcripts. One cloned sequence corresponded to the canonical *PLIN3A* coding sequence (CDS) (*PLIN3-201*, UniProt ID: <u>O60664-1</u>, CCDS12137, transcript ID <u>ENST00000221957.9</u>). A second cDNA encoded a 126 bp, 42 AA-shorter sequence, corresponding to an alternative splicing event in *PLIN3A* exon 6, with the Chr19:4844794-4847816 junction being used over Chr19:4844794-4847690 (figure 1A-C). The shorter CDS, *PLIN3B*, was not reported in Ensembl (Dyer et al., 2025). ENSE00002796125, the exon generated by the alternative splicing event was however present in the <u>ENST00000589163.5</u> transcript (*PLIN3-205*, UniProt ID: <u>K7ERZ3</u>), which was manually annotated by the HAVANA (Human and Vertebrate Analysis and Annotation) group as a 5’ incomplete CDS (figure 1D-E). Having cloned the full CDS, we propose that the *PLIN3B* CDS corresponds to the full sequence of PLIN3-205 (figure 1E).

**Figure 1.**
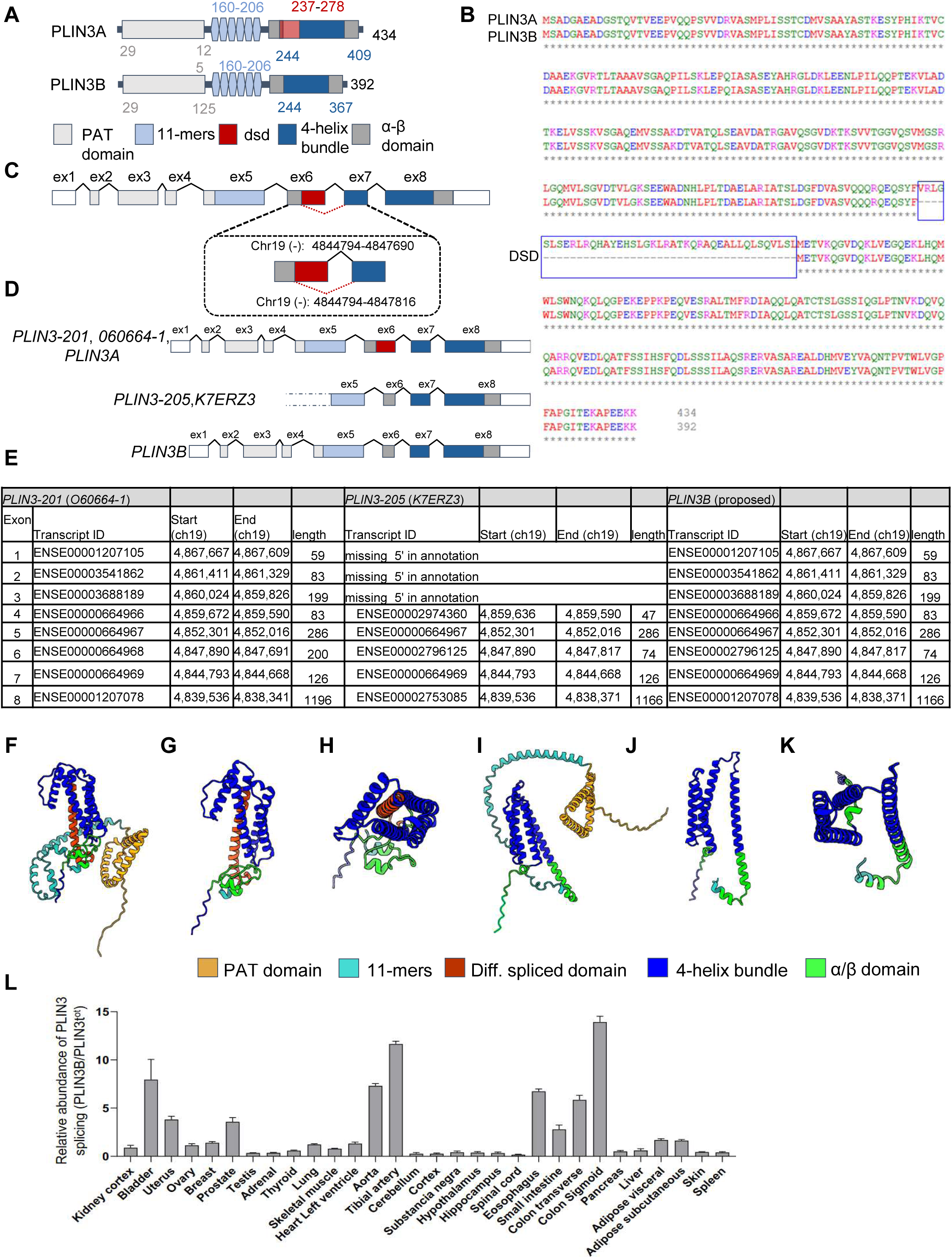
PLIN3B is a minor PLIN3 splicing variant. **A)** Schematics of human PLIN3A and PLIN3B structures and their domains. Colored numbers indicate the initial and final residues of the respective domains. DSD=differentially spliced domain. **B)** Amino acidic sequence of PLIN3A and PLIN3B. Blue square indicates the DSD. **C)** Schematics of the splicing event generating *PLIN3B*. **D)** Exon structure of *PLIN3A*, *PLIN3B* and of the *PLIN3-205/K7ERZ3* accession. Color code corresponding to protein domains as in figure 1A. **E)** Table of individual Ensembl transcripts of *PLIN3A*, *PLIN3B* and *PLIN3-205/K7ERZ3*. **F-K)** AlphaFold2 prediction of PLIN3A and PLIN3B. From left to right: PLIN3A, PLIN3A (206-434) side view, PLIN3A (206-434) top view, PLIN3B, PLIN3B (206-392) side view, PLIN3B (206-392) top view. **L)** *PLIN3B* expression pattern in human tissues from GTEx database.

The PLIN3B 42 AA differentially spliced domain (DSD) encompasses a portion of one of the four α-helixes in the four-helix bundle (4HB) domain (figure 1A). We predicted the PLIN3B structure with AlphaFold (Jumper et al., 2021), which showed a partial preservation of the 4HB domain structure, with a new α-helix formed by the N-terminal residues of the α/β domain (PLIN3B:222-266). Consequently, the α/β domain (which in PLIN3A is composed by residues located at the N- and C-terminal of the 4HB domain) (Hickenbottom et al., 2004), appears to be disrupted and the two moieties separated (figure 1F-K, supp. figure 1A-D). In the GTEx database (Lonsdale et al., 2013), *PLIN3B* appears to be a minor splicing variant, with the highest relative expression of the ENSE00002796125 exon detected particularly in smooth muscle-rich tissues such as in the vasculature and the gastrointestinal tract (figure 1L).

### PLIN3B localizes to mitochondria in addition to LD

We cloned *PLIN3A* and *PLIN3B* into P2A self-cleaving plasmids, allowing coexpression of the CDS and blue fluorescent protein (BFP) (Kim et al., 2011). We transfected HeLa cells with the two plasmids and immunostained with an antibody targeting the PLIN3 11-mers domain (residues 73-225) (table 1), which is conserved in both isoforms (figure 1A). While PLIN3A diffusely stained the cytosol, PLIN3B localized to mitochondria in cells cultivated both in standard DMEM (supp. figure 2A) and in DMEM supplemented with oleic acid (OA) to stimulate LD biogenesis (figure 1G). We confirmed the mitochondrial localization by performing subcellular fractionation of PLIN3A-P2A-BFP and PLIN3B-P2A-BFP transfected cells, highlighting an enrichment of PLIN3B in the mitochondrial fraction (figure 2B-D). We additionally cloned PLIN3A and PLIN3B into an N-terminal 3×FLAG plasmid. Immunofluorescence of FLAG-PLIN3B- and FLAG-PLIN3A-transfected HeLa cells confirmed PLIN3B mitochondrial localization (supp. figure 1H). Both FLAG-PLIN3A and FLAG-PLIN3B displayed additional LD staining (supp. figure 1H), providing independent validation of localization.

**Figure 2.**
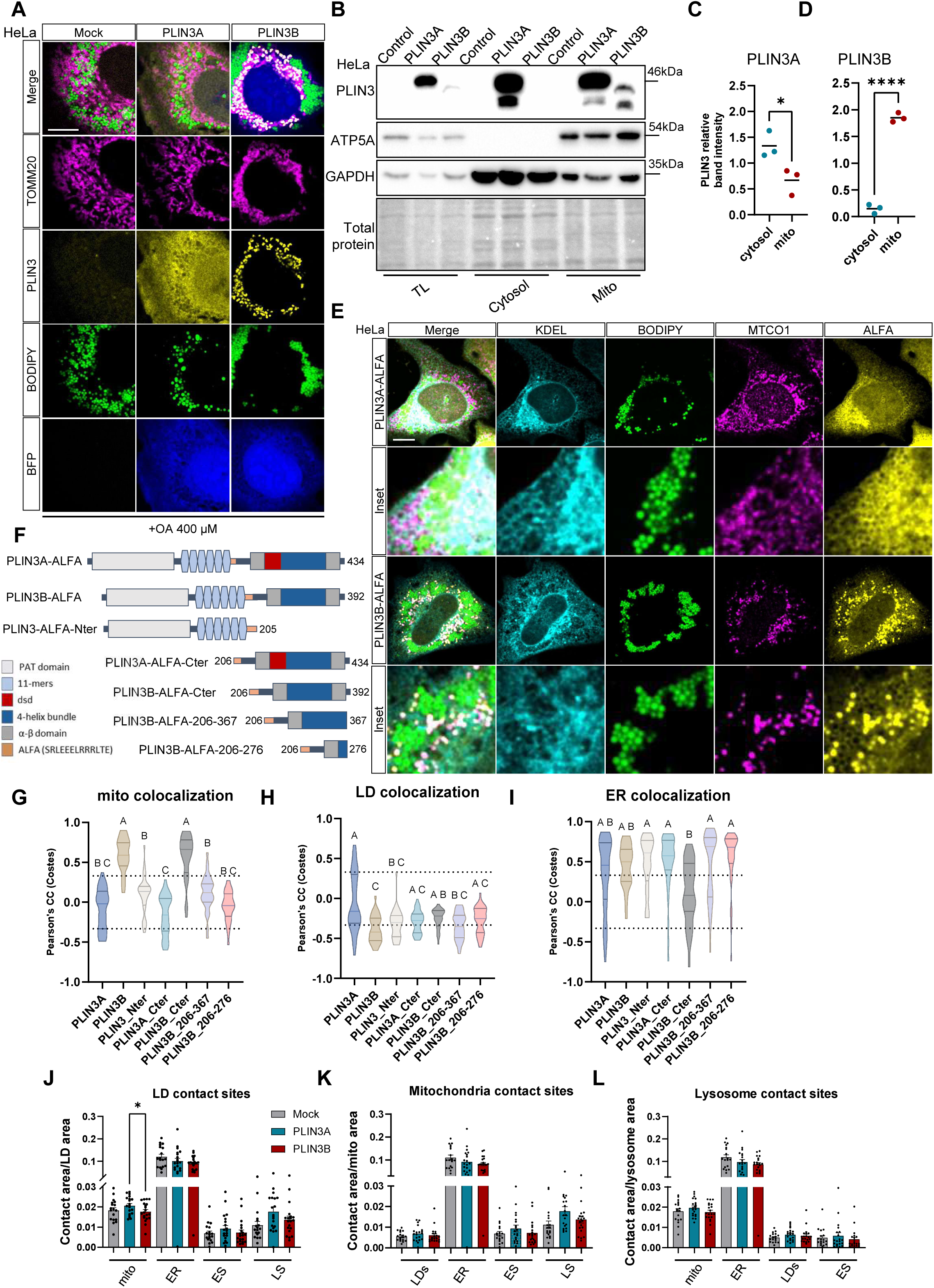
PLIN3B localizes at mitochondria. **A)** Representative immunocytochemistry confocal images of HeLa cells transfected with PLIN3A-P2A-BFP or PLIN3B-P2A-BFP plasmids and incubated with OA. Cells were stained with PLIN3 and TOMM20 (mitochondria) antibodies and with BODIPY 493/503 (LD). Images were acquired with LSM780. Scale bar=10 µm. N=3 independent experiments. **B)** WB of cellular fraction of HeLa cells transfected with PLIN3A-P2A-BFP, PLIN3B-P2ABFP or with the control plasmid FLAG-P2A-BFP. Expression of PLIN3, ATP5A and GAPDH was assayed. N=3 independent experiments. **C)** Quantification of PLIN3A band intensity in cytosolic and mitochondrial fractions. Individual band intensity value was normalized by mean band intensity for every experiment, Error bar=mean, ∗*p* < 0.05, ^∗∗^*p* < 0.01 (unpaired *t* test), N=3 independent experiments. **D)** Quantification of PLIN3B band intensity in cytosolic and mitochondrial fractions. Individual band intensity value was normalized by mean band intensity for every experiment, error bar=mean, ∗*p* < 0.05, ^∗∗^*p* < 0.01 (unpaired *t* test), N=3 independent experiments. **E)** Representative immunocytochemistry confocal images of HeLa cells transfected with different PLIN3A-ALFA and PLIN3B-ALFA and incubated with OA. Cells were stained with antibodies against ALFA, MTCO1 (mitochondria), KDEL (ER), and with BODIPY 493/503 (LD). Images were acquired with LSM780. Scale bar=10 µm. N=3 independent experiments. **F)** Schematics of ALFA tagged PLIN3A and PLIN3B constructs used in **figure 2E, supplemental figure1I**. **G-I)** Quantification of Pearson’s correlation coefficient of ALFA and MTCO1 **(G)**, ALFA and Bodipy 493/503 **(H)** and ALFA and KDEL **(I)** cytosolic signals. Kruskal-Wallis test. Violin plot lines represent median and quartiles. Distinct letters (A, B, C) indicate significant differences between groups. n=31-44 cells per condition from 3 independent experiments. **2J-L)** quantification of LD **(J)**, mitochondria **(K)**, and lysosome **(L)** contact sites with other organelles in mock, PLIN3A-P2A-BFP and PLIN3B-P2A-BFP-transfected cells in multispectral imaging experiment. Error bar=mean + SEM. Mixed-effects analysis, with the Geisser-Greenhouse correction, followed by two-stage linear step-up procedure of Benjamini, Krieger and Yekutieli with individual variances computed for each comparison. Mito=mitochondria, LS=lysosomes, ES=endosomes. N=18-20 cells per condition from 2 independent experiments.

**Table 1:**
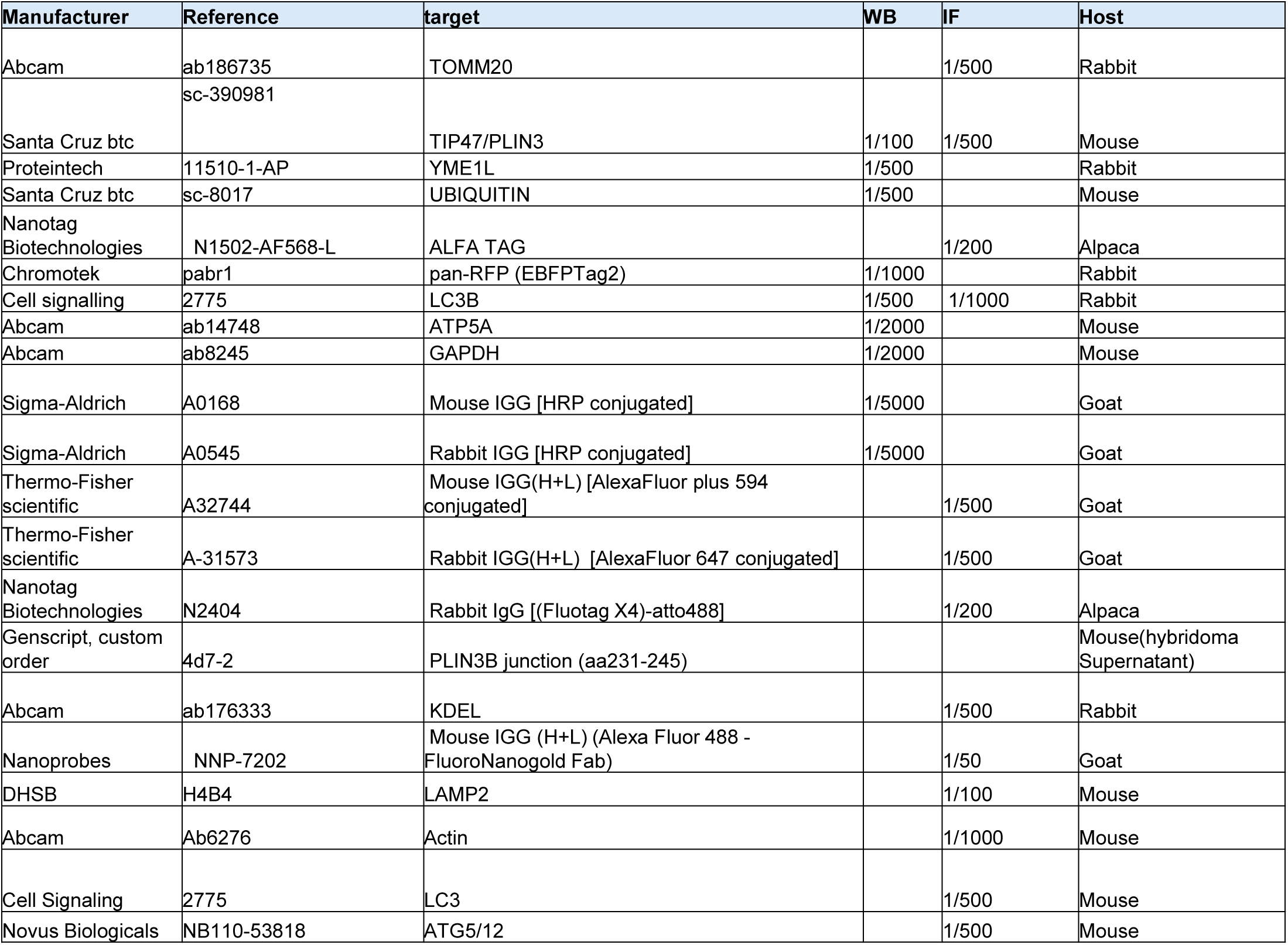
list of antibodies. Antibodies used in Immunofluorescence and western blot in the manuscript.

### PLIN3 C-terminal residues are required for PLIN3B mitochondrial localization

In order to confirm the link between PLIN3B differential splicing and organelle targeting, we generated full-length PLIN3A and PLIN3B tagged with the ALFA peptide tag (Gotzke et al., 2019), and multiple ALFA-tagged fragments of the two proteins (figure 2E-F, supp. figure 1I). The 15 AA-long tag was inserted at the end of the 11-mers domain (Proline 205) in all the constructs to avoid differential positional effects. AlphaFold prediction did not highlight notable alterations of the 4HB domain caused by insertion of the tag in PLIN3A or PLIN3B (supp. figure 1C-F).

We co-stained the PLIN3 constructs with ER, mitochondria and LD markers. PLIN3A-ALFA and PLIN3B-ALFA displayed similar targeting to FLAG-PLIN3A and FLAG-PLIN3B. PLIN3A-ALFA localized at LDs and diffusely stained the cytosol, while PLIN3B predominantly localized at mitochondria, with only minor staining of LDs. To quantify the organelle targeting of the PLIN3 fragments, we performed colocalization analysis of PLIN3-ALFA fragments and the different organelles (figure 2G-I).

PLIN3B-ALFA and PLIN3B-ALFA-CTER consistently colocalized with mitochondria. PLIN3B-ALFA-206-367 and PLIN3B-ALFA-206-276 did not accumulate at mitochondria, suggesting that residues conserved in both isoforms are responsible for PLIN3B mitochondrial targeting (figure 2G). Full-length PLIN3A-ALFA was the only construct to display positive correlation of ALFA and LD signals (figure 2H).

PLIN3 is known to accumulate at the ER and to mediate LD biogenesis (Bulankina et al., 2009). PLIN3A-P2A-BFP, but not PLIN3B-P2A-BFP, increased LD formation in OA treated cells (supp. figure 2A). To exclude PLIN3B colocalization with mitochondria as a consequence of PLIN3B accumulation at the ER, we analyzed ALFA and ER colocalization of all PLIN3 constructs. None of the constructs showed a preferential colocalization with the ER (figure 2I).

### PLIN3B mitochondrial localization is independent from organelle contacts

To exclude PLIN3B mitochondrial localization to be dependent from the involvement of other organelles, we analyzed the effect of PLIN3B on organelles and organelle contacts with multispectral imaging (Cohen et al., 2018), allowing to simultaneously analyze LD, ER, mitochondria, endosome and lysosomes morphology and interactions.

Despite colocalizing with mitochondria, PLIN3B transfected cells displayed reduced LD-mitochondria contacts confronted to PLIN3A (Figure 2J) and showed no differences in organelles contacts involving ER, endosomes and lysosomes (Figure 2K-M). PLIN3B increased mitochondrial area, but showed no effect on LD, ER, endosome and lysosome area (supp. Figure 2B). Together these results suggest that PLIN3B localization at mitochondria is independent from LD, ER, endosomes and lysosomes.

### PLIN3B mitochondrial localization is independent of autophagolysosome-associated pathways

PLIN3 is known to be targeted to lysosomes through its KFERQ motif by interaction with the heat-shock cognate 71 kDa protein (HSC70), which targets the protein to lysosomes through LAMP2 (Kaushik and Cuervo, 2015). Given the multiple mitochondrial quality control systems involving interaction between mitochondria and the autophagolysosomal pathway, we wanted to exclude PLIN3B mitochondrial localization being secondary to organelle contacts. In HeLa cells, PLIN3B-ALFA did not colocalize with LAMP2, with or without OA (supp. figure 2C-D).

PLIN3B mitochondrial localization was unaffected in autophagy-impaired ATG5 KO cells (Wang et al., 2008) (supp. figure 2E-F). Thus, using independent approaches, we excluded PLIN3B mitochondrial localization to autophagosome formation. This distinguishes canonical PLIN3A lysosomal targeting from PLIN3B mitochondrial localization, which does not depend on autophagy.

### PLIN3B is localized inside mitochondria

Having excluded PLIN3B targeting to other organelles as the reason for the mitochondrial staining observed, we focused on the sub-mitochondrial localization of the new isoform. Mitochondria are compartmentalized organelles with defined import mechanisms that regulate protein delivery to the outer mitochondrial membrane (MOM), inner mitochondrial membrane (MIM), intermembrane space (IMS), and matrix. PLIN3 does not feature a mitochondrial targeting sequence (Rath et al., 2021), and consistent with this, PLIN3B was not predicted as mitochondrial using Deepmito (Savojardo et al., 2020). PLIN1 and PLIN5 are known to associate with the MOM by interacting with mitochondrial proteins at LD-mitochondria contacts (Boutant et al., 2017; Miner et al., 2023). Despite a report of PLIN3 association with mitochondria in the literature, likely reflecting increased LD-mitochondria proximity after training (Ramos et al., 2015), PLIN3B did not induce tethering of the two organelles in our experiments (figure 2A, E, H). In electron micrographs, PLIN3B-transfected cells displayed swollen mitochondria, with alterations of the cristae network and abnormal distribution of IMS and matrix (figure 3A, supp. figure 3B). PLIN3A-transfected cells and control cells did not show these alterations (figure 3B, supp. figure 3A). Despite the morphological alterations, mitochondria in PLIN3B-transfected cells retained polarization-dependent labeling by MitoTracker Red CMXRos (Neikirk et al., 2023), suggesting preservation of mitochondrial membrane potential (supp. figure 3C).

**Figure 3.**
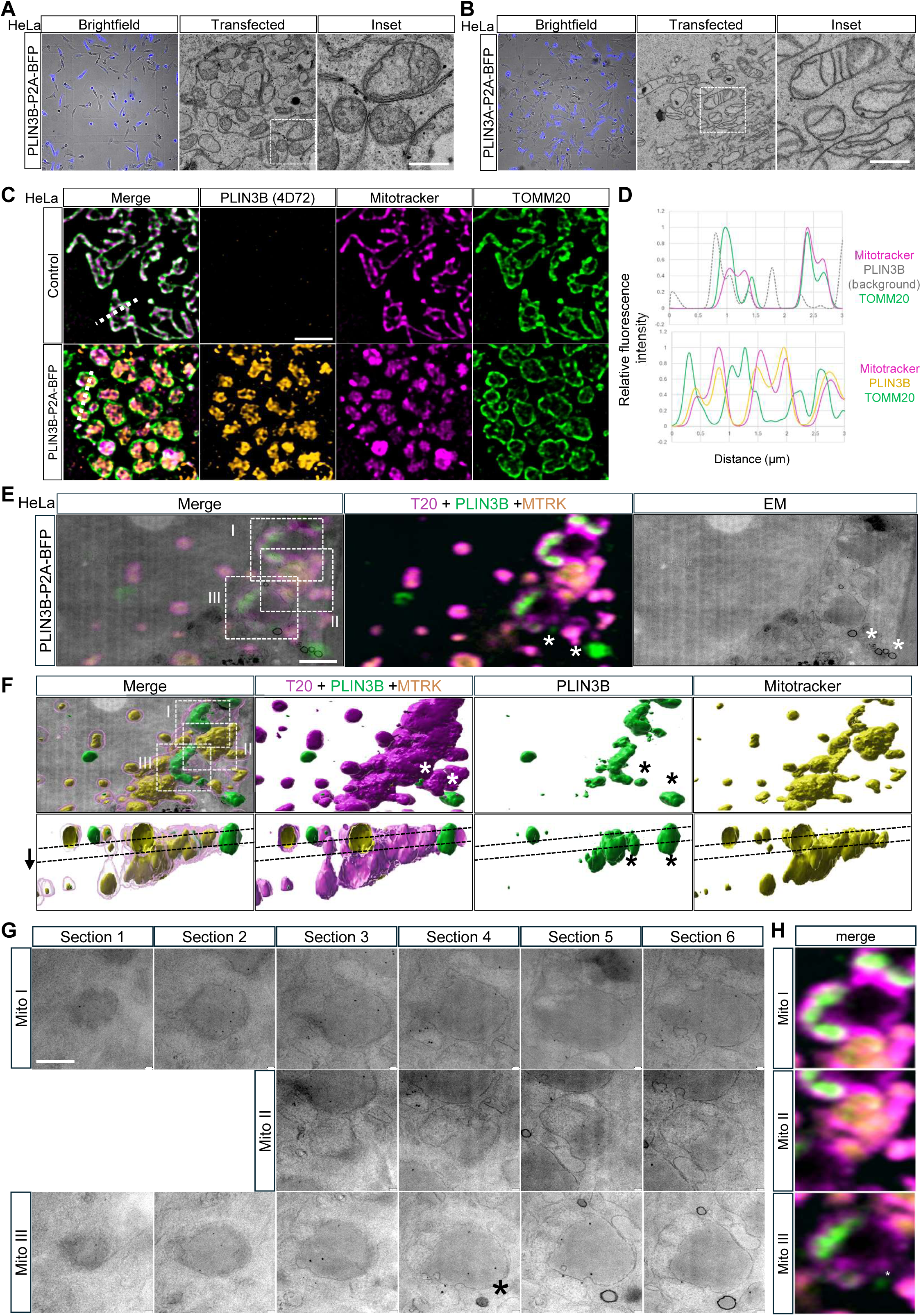
PLIN3B is localized inside mitochondria. **A)** Representative electron micrographs of cells transfected with PLIN3B-P2A-BFP. Scale bar=500 nm. **B)** Representative electron micrographs of cells transfected with PLIN3A-P2A-BFP. Scale bar=500 nm. **C)** Representative immunocytochemistry structured illumination microscopy (SIM^2^) images of mock transfected HeLa cells or HeLa cells transfected with PLIN3B-P2A-BFP. Cells were stained with MitoTracker Red CMXRos, anti-PLIN3B 4D72 antibody and with anti-TOMM20 antibody. Images were acquired with Elyra7 microscope. Scale bar=2.5 µm. N=3 independent experiments. **D)** Profile plot of dashed lines in **figure 3C**. **E)** Correlation between electron micrograph and CLSM images of HeLa cells transfected with PLIN3B-P2A-BFB. The images correspond to the ROI in **supplemental figure 3E**. White asterisk correspond to the same structure as in **figure 3F-H**. Dashed ROIs corresponding to serial sections in **figure 3H**. Scale bar=2 µm. **F)** Imaris 3D reconstruction of the ROI in **supplemental figure 3E** in top and side view. Black dashed lines indicate the top and bottom serial sections in **figure 3H**. Black arrow indicates serial sections direction. White and black asterisk correspond to the same structure as in **figure 3E** and **3G-H**. **G)** Electron micrographs of serial sections from mitochondria in ROIs from **figure 3E**. White asterisk corresponds to the same structure as in **figure 3E-F** and **3H**. Scale bar=1 µm. **H)** CLSM image corresponding to section 4 in **figure 3G**. White asterisk corresponds to the same structure as in **figure 3E-G**. Cells were stained with MitoTracker Red CMXRos, PLIN3B 4D72 antibody and with anti-TOMM20 antibody.

To exclusively detect PLIN3B in transfected cells, we generated a custom isoform-specific antibody targeting the new splicing junction, called 4D72 (supp. figure 5A-C). We obtained structured illumination microcopy (SIM^2^) super-resolved images of PLIN3B-P2A-BFP transfected cells stained with TOMM20 antibody (labeling MOM) and MitoTracker Red CMXRos (labeling MIM and matrix), together with the custom PLIN3B antibody. Mitochondria of PLIN3B-transfected cells were round and enlarged compared with control cells (figure 3C), consistent with our observations in electron micrographs. PLIN3B localized inside the TOMM20-stained MOM and colocalized with MitoTracker Red CMXRos (figure 3C-D).

To limit misinterpretations of the light microscopy results due to limitations in axial resolution, we developed a fluoronanogold-correlative light-electron microscopy (FNG-CLEM) protocol (supp. figure 3D). Detergent-free liquid nitrogen permeabilization enabled multiplex immunocytochemistry while preserving cellular membrane integrity (figure 3E-H) (Knott et al., 2009). We performed three-dimensional imaging of PLIN3B-P2A-BFP-transfected cells and 3D reconstruction of PLIN3B, TOMM20 and MitoTracker signals with Imaris (supp. figure 3E). The same sample was silver-enhanced and processed to obtain serial ultrathin sections of a portion of the cell volume (figure 3F, black dashed lines). Light and electron microscopy images were correlated based on cell, nucleus and mitochondrial contour as well as PLIN3B signal around LDs (figure 3E-H, black and white asterisks), which were unambiguously detectable in both light microscopy stacks and serial sections of silver-enhanced nanogold particles on fluoronanogold secondary antibody detected by transmission electron microscopy (ssTEM). MitoTracker and PLIN3B signals were contained inside mitochondria in the 3D reconstruction (figure 3F). Silver-enhanced nanogold signal was observed inside mitochondria in multiple serial sections TEM (figure 3G). CLEM was additionally performed in non-silver-enhanced samples, showing similar localization of PLIN3B signal inside mitochondria by correlating the fluorescent signal of the secondary antibody by CLSM and the ultrastructure on serial sections TEM using the cell contours, the nuclei and the labeling of the mitochondria as registration cues (supp. Figure 3F).

### PLIN3B co-purifies with proteasome subunits and mitochondrial proteins

We performed PLIN3B coimmunoprecipitation and mass spectrometry analysis (coIP-MS) in 3×FLAG-PLIN3B-expressing cells and identified several PLIN3B interactors (figure 4A). Gene ontology analysis of the interactors highlighted an enrichment of proteasome subunits (figure 4B-C). Additionally, multiple Gene Ontology Cell Compartment (GOCC) groups were assigned to mitochondrial proteins. We cross-checked the list of proteins enriched in FLAG-PLIN3B coimmunoprecipitates with the Mitocarta3.0 database (Rath et al., 2021). This resulted in 25 putative mitochondrial interactors, with a predominance of MIM and IMS proteins (figure 4D). These results support the intra-mitochondrial localization of PLIN3B suggested by super-resolution imaging and confirmed by the FNG-CLEM approach.

**Figure 4.**
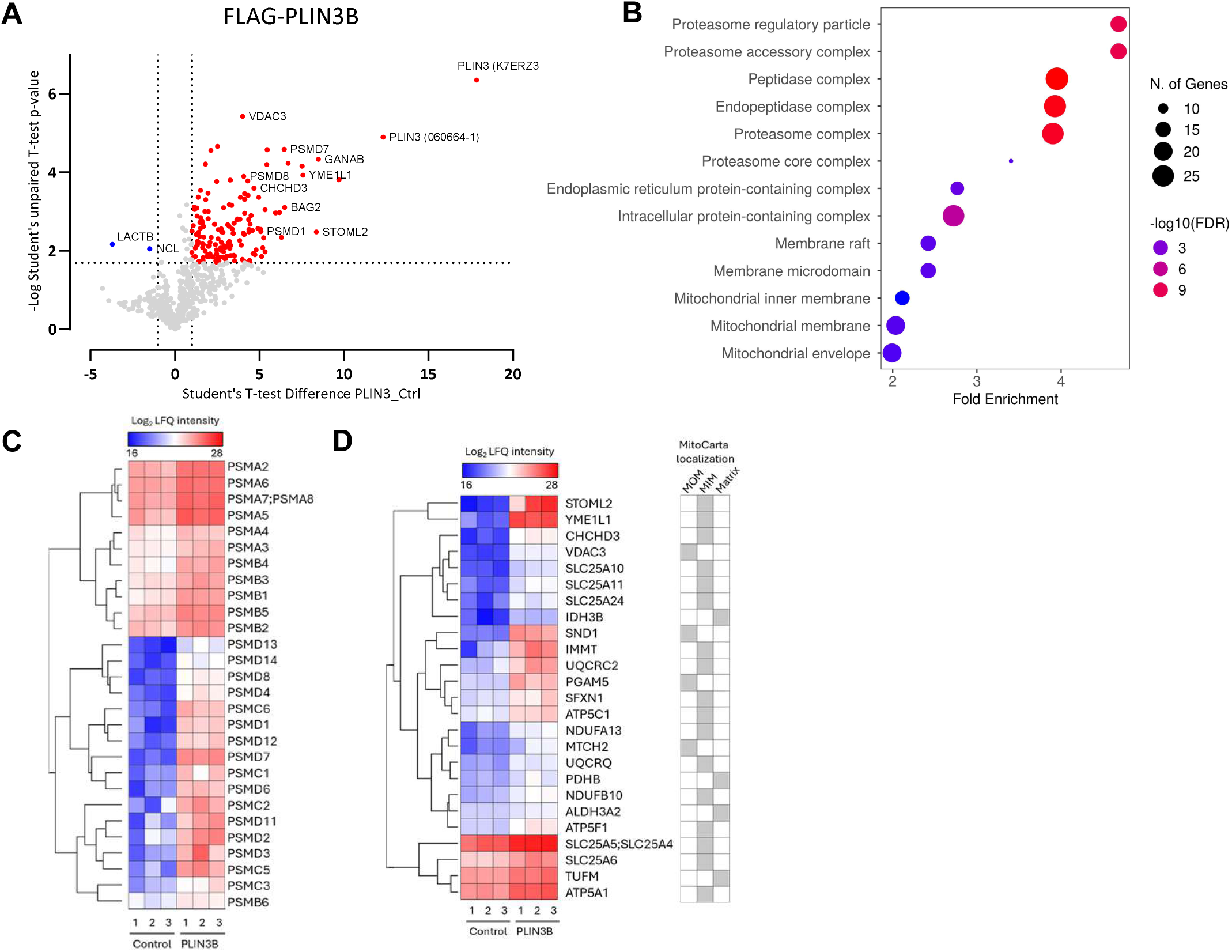
PLIN3B co-purifies with mitochondrial and proteasome proteins. **A)** Volcano plot of protein detected in FLAG-PLIN3B co-IP/MS analysis. Proteins significantly enriched in FLAG-PLIN3B sample are highlighted in blue, while proteins significantly enriched in MOCK sample are highlighted in red. N=3 independent experiments, *p* < 0.05 (*t* test with multiple testing correction: permutation-based FDR at 0.05). **B)** GOCC analysis of proteins significantly enriched in 3xFLAG-PLIN3B co-IP/MS analysis. **C)** Heatmap of Log_2_ LFQ intensity of proteins enriched in FLAG-PLIN3B samples part of the Gene Ontology Cell Compartment (GOCC) “proteasome complex” in gene ontology enrichment analysis. Hierarchical clustering was performed with 1-Pearson’s correlation method. **D)** Heatmap of Log_2_ LFQ intensity of proteins enriched in FLAG-PLIN3B samples and labelled as mitochondrial in the MitoCarta3.0 database. On the right, the MitoCarta3.0-attributed compartment of each protein. Hierarchical clustering was performed with 1-Pearson’s correlation method.

### Antisense oligonucleotide treatment confirms PLIN3B splicing event

To confirm that PLIN3B is generated by the proposed alternative splicing event, we designed a morpholino antisense oligonucleotide (AON) targeting the exon 6 canonical splicing site (table 2). We treated HeLa cells with increasing concentrations of *PLIN3A*-targeting AON or a scramble control AON and performed retrotranscription-endpoint-PCR (RT-epPCR) to detect variations in the abundance of *PLIN3A* and *PLIN3B* mRNA. Endo-Porter-treated cells displayed a single band corresponding to the predicted *PLIN3A* amplicon (figure 5A). In AON-treated cells, an additional shorter band was observed, consistent with the amplicon originating from *PLIN3B* cDNA amplification (figure 5A-C). Sequencing of the shorter amplicon confirmed the presence of the *PLIN3B* splicing junction (supp. figure 4A). A second PCR primer pair targeting the splicing junction, gave similar results (figure 5A, supp. figure 4B). Using the primer pair employed for Gateway cloning, a third PCR assay revealed a shorter band after *PLIN3A* AON treatment, corresponding to the full *PLIN3B* cDNA (figure 5A, supp. figure 4C). We quantified the abundance of *PLIN3A* and *PLIN3B* mRNA by qPCR, highlighting a more than 100-fold increase in *PLIN3B* mRNA after AON treatment (supp. figure 4D). Consistently, *PLIN3A* levels decreased and overall *PLIN3* levels remained stable.

**Table 2:**
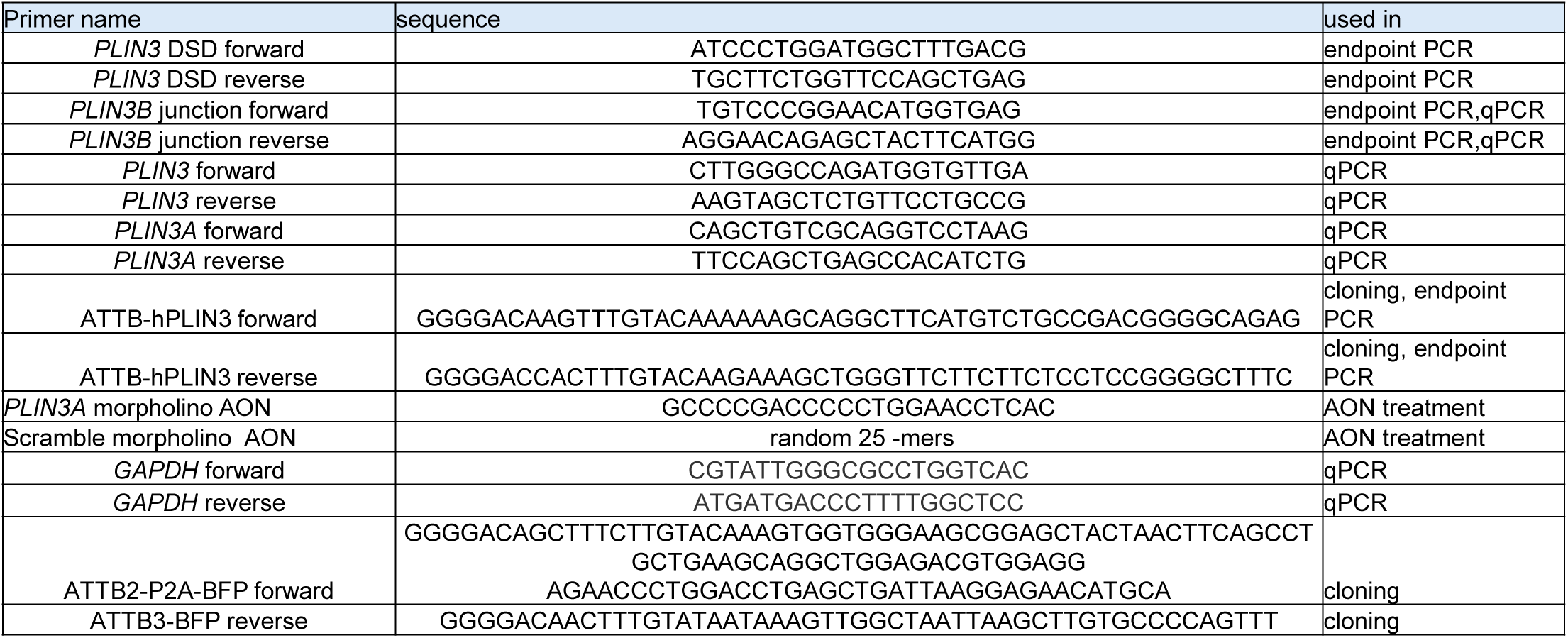
list of primers and oligonucleotides. Primers and oligonucleotides used in cloning, PCR and AON treatment.

**Figure 5.**
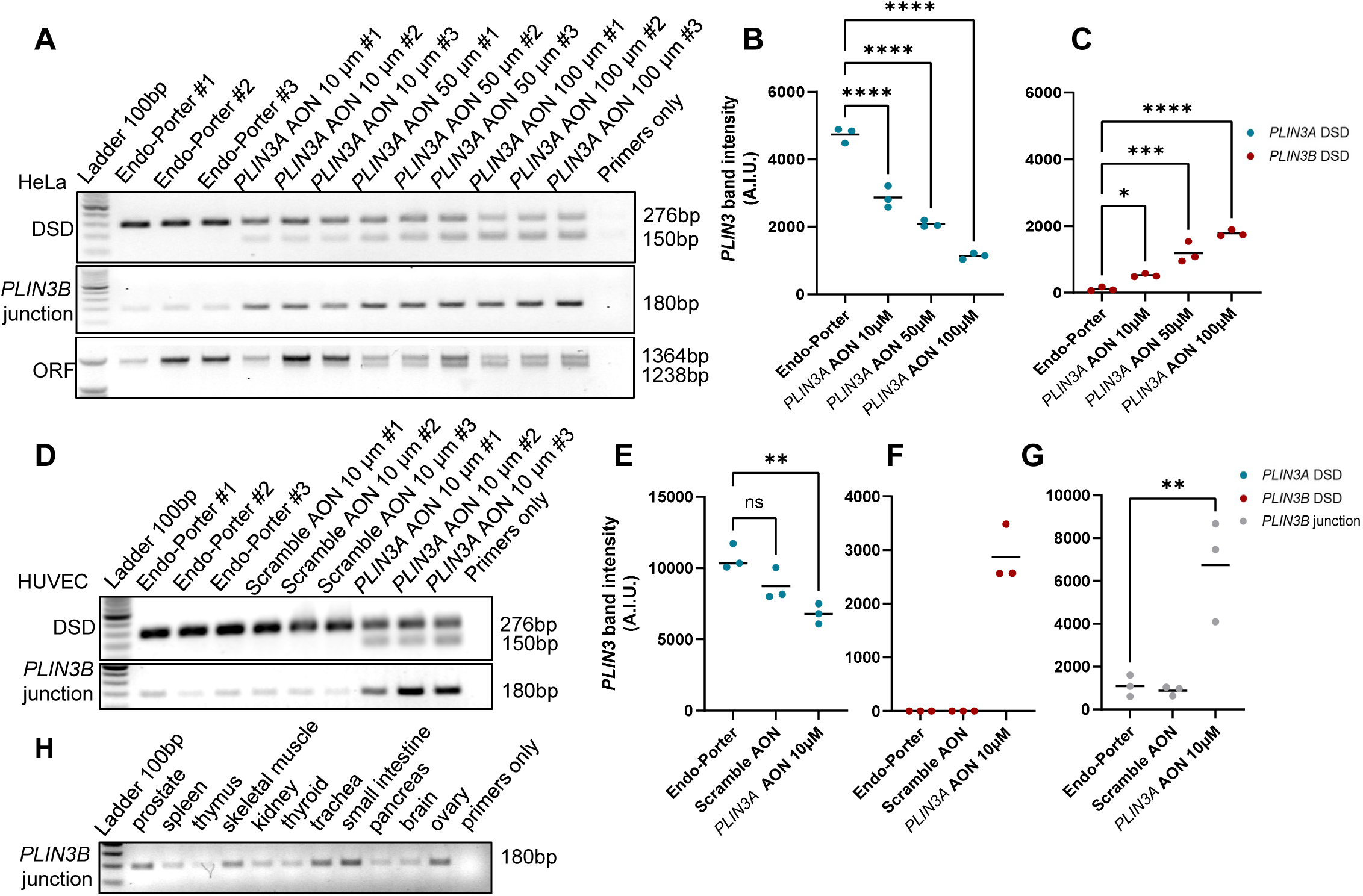
Detection and antisense oligonucleotide modulation of *PLIN3B* splicing event. **A)** Agarose gel-PCR of cDNA from antisense oligonucleotide treated HeLa cells. Primers were designed to target the DSD region, the *PLIN3B* junction and the whole ORF of *PLIN3A* and *PLIN3B* transcripts. N=3 independent experiments. **B-C)** Quantification of *PLIN3A* **(B)** and *PLIN3B* **(C)** DSD amplicon band intensity from (A). ∗∗∗∗p < 0.0001, ∗∗∗p < 0.001, ∗∗p < 0.01, ∗p < 0.05 (Ordinary one-way ANOVA), N=3 independent experiments. **D)** Agarose gel-PCR of cDNA from AON treated HUVEC cells. Primers were used targeting DSD region, the *PLIN3B* junction and the whole ORF of *PLIN3A* and *PLIN3B* transcripts. N=3 independent experiments. **E-G)** Quantification of *PLIN3A* DSD amplicon **(E)**, *PLIN3B* DSD amplicon **(F)** and *PLIN3B* junction amplicon **(G)** band intensity from (D). ∗∗∗∗p < 0.0001, ∗∗∗p < 0.001, ∗∗p < 0.01, ∗p < 0.05 (Ordinary one-way ANOVA), N=3 independent experiments. **H)** Agarose gel-PCR of cDNA from Ambion FirstChoice RNA panel. Primers were selected to target the *PLIN3B* junction.

We confirmed morpholino-induced splicing modulation in human umbilical vein endothelial cells (HUVEC), a non-cancerous, non-immortalized human cell line (figure 5D-G).

We next employed the *PLIN3B* junction PCR assay to investigate the presence of the *PLIN3B* splicing event in human tissues. PCR performed on an Ambion FirstChoice human RNA survey panel revealed the differential presence of *PLIN3B* junction across human tissues (figure 5H), with partial overlap with GTEx expression data (figure 1L).

### PLIN3B exhibits reduced stability compared to PLIN3A

Overexpression of PLIN3A and PLIN3B in cell fractionation and in PLIN3B antibody validation experiments revealed differences in the expression levels of the two isoforms (figure 2B, supp. figure 5C-D). Given that colon had high levels of *PLIN3B* junction expression (figure 1L), we used HCT116 colon cancer cells for the 4D72 antibody validation. Overexpressed PLIN3B was approximately 30-fold less abundant than overexpressed PLIN3A, and its expression levels were comparable to endogenous PLIN3A (supp. figure 5D). This observation suggested that PLIN3B may have a shorter half-life than PLIN3A. We then performed cycloheximide (CHX) pulse-chase on PLIN3A-P2A-BFP and PLIN3B-P2A-BFP transfected cells (figure 6A-C, supp. figure 5E-F). While PLIN3A/BFP levels remained stable, the PLIN3B/BFP ratio displayed a rapid decay after an initial spike at 15 min, with an estimated half-life of approximately 1 h.

**Figure 6.**
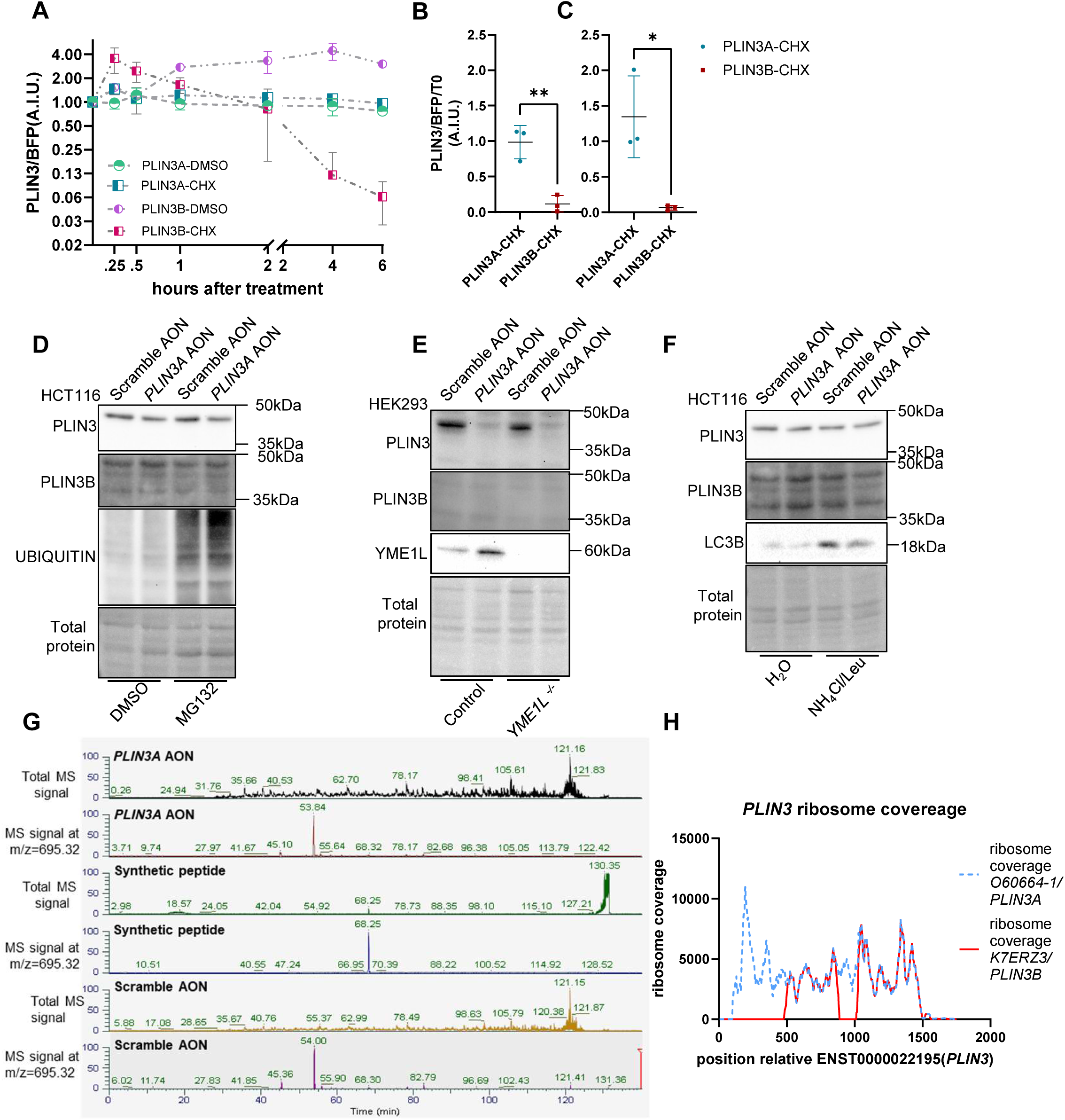
PLIN3B endogenous protein expression is non-detectable. **A)** PLIN3A and PLIN3B levels in Cycloheximide (CHX) pulse-chase in PLIN3A-P2A-BFP and PLIN3B-P2A-BFP transfected HCT116 *PLIN3^-/-^*cells. N=3 independent experiments. **B-C)** Comparison of BFP normalized PLIN3A and PLIN3B levels at 4h (left) and 6h (right) in cycloheximide treated HCT116 *PLIN3^-/-^* cells. ∗∗p < 0.01, ∗p < 0.05 (unpaired t-test). N=3 independent experiments. **D)** Representative WB of Scramble AON (50 μM), *PLIN3A* AON (50 μM) treated HCT116 cells incubated with the proteasome inhibitor MG132 (5 μM). Expression of PLIN3, PLIN3B and total UBIQUITIN was assayed. N=3 independent experiments. **E)** Representative WB of Scramble AON (100 μM), *PLIN3A* AON (100 μM) treated *WT* and *YME1L^-/-^* HEK293 cells. Expression of PLIN3, PLIN3B and YME1L was assayed. N=3 independent experiments. **F)** Representative WB of Scramble AON (50 μM), *PLIN3A* of *PLIN3A* AON (50 μM) treated HCT116 cells incubated with the lysosome inhibitors NH_4_Cl (20 mM) and leupeptin (Leu) (100 μM). Expression of PLIN3, PLIN3B and LC3B was assayed. N=3 independent experiments. **G)** Total Ion Chromatogram traces of Scramble AON (50 μM), *PLIN3A* AON (50 μM) treated HCT116 cells samples and synthetic peptide standard. **H)** *PLIN3A* and *PLIN3B/K7ERZ3* ribosome coverage from Trips-Viz transcriptome browser.

### PLIN3B protein remains below detection limits in steady-state and protease-defective cells

AON treatment of HCT116 cells induced detectable *PLIN3B* mRNA levels (supp. figure 5G) and reduced PLIN3A protein levels (figure 6D, supp. figure 5H). However, PLIN3B protein was not detectable using either the PLIN3B-specific custom-made antibody or non-specific PLIN3 antibody (figure 6D).

Given that mass spectrometry analysis revealed enrichment of proteasome complex subunits among PLIN3B interactors (figure 4A-C), we hypothesized that PLIN3B protein remaining undetectable was due to proteasome-mediated degradation, as previously reported for the closely related PLIN2 (Takahashi et al., 2016). Lower PLIN3B expression in transfection experiments and its shorter half-life compared to PLIN3A supported this hypothesis (figure 6A-C, supp. figure 5D). We inhibited proteasomal activity with MG132 (Han et al., 2009). MG132 treatment increased the level of total ubiquitinated proteins but did not affect PLIN3A protein levels and did not enable PLIN3B detection (figure 6D).

We next examined the potential involvement of the mitochondrial protease YME1-like protein 1 (YME1L). YME1L is an IMS protein with reported chaperone-protease activity (MacVicar et al., 2019; Stiburek et al., 2012). Basal YME1L expression was shown to maintain the concentration of its mitochondrial proteolytic substrates below detection levels (Wai et al., 2016). STOML2, a YME1L interactor involved in proteostasis regulation (Wai et al., 2016), was also identified among PLIN3B interactors in our CoIP-MS experiment (figure 4A-B, 4D). We assessed the effect of YME1L loss on PLIN3B detection in AON-treated cells. Despite reducing PLIN3 levels (supp. figure 5I), AON treatment did not enable PLIN3B detection in both HEK293 wild-type or YME1L knockout cells (figure 6E).

A third proposed mechanism for PLIN3B rapid turnover was the degradation by the autophagosome-lysosome pathway. PLIN3 is known to be targeted to lysosomes and degraded via HSC70 interaction (Kaushik and Cuervo, 2015). We inhibited lysosomal acidification, and consequently lysosome-mediated protein degradation, with a combination of leupeptin and ammonium chloride (NH_4_Cl) (Seglen et al., 1979). Despite a borderline reduction of PLIN3A (supp. figure 5J), NH_4_Cl plus leupeptin treatment did not enable detection of PLIN3B (figure 6F).

### PLIN3B is not detected by targeted proteomics in morpholino-treated cells

We performed targeted mass spectrometry analysis of scramble AON- and PLIN3A AON-treated cells to assess PLIN3B protein detection independently of antibody-based approaches. The K7ERZ3/PLIN3B junction generates a unique tryptic peptide, QEQSYFMETV, which was clearly detectable in the FLAG-PLIN3B coIP-MS analysis (figure 4A, table 3). As an additional confirmation of PLIN3B detectability, the purified immunization peptide QEQSYFMETVKQGVDC was used as a positive control. Tryptic digestion of this peptide generated a clear peak at m/z 695.32 (2+ ion), corresponding to the splicing junction peptide QEQSYFMETV (figure 6G).

**Table 3:**
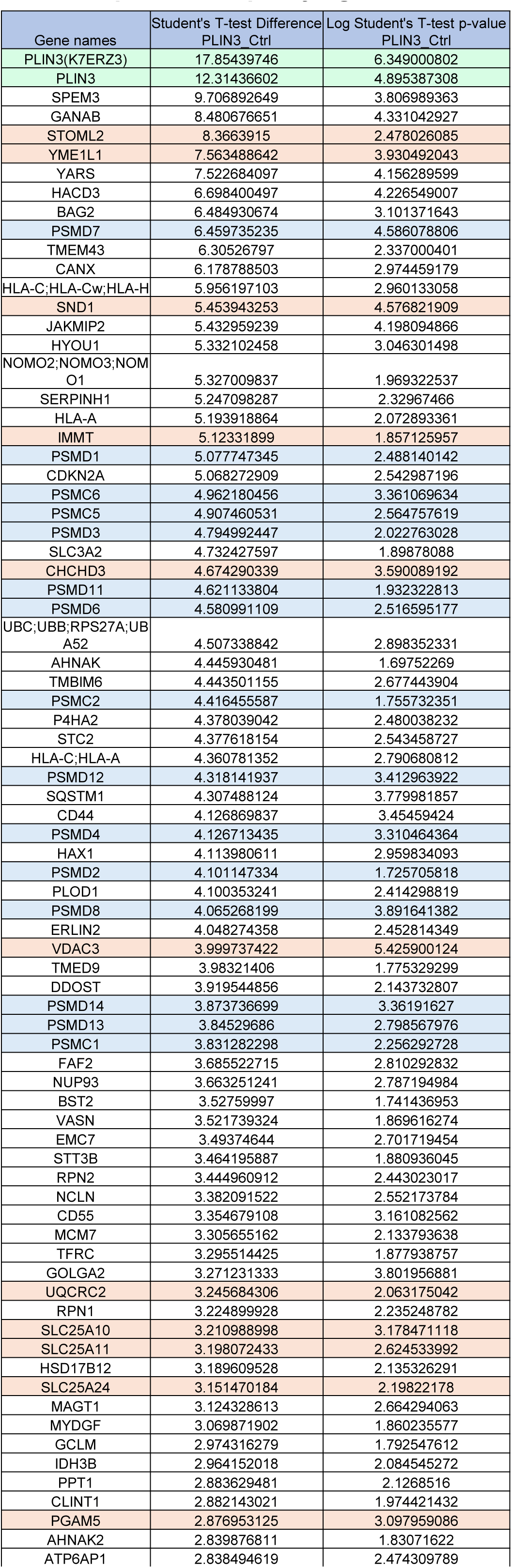
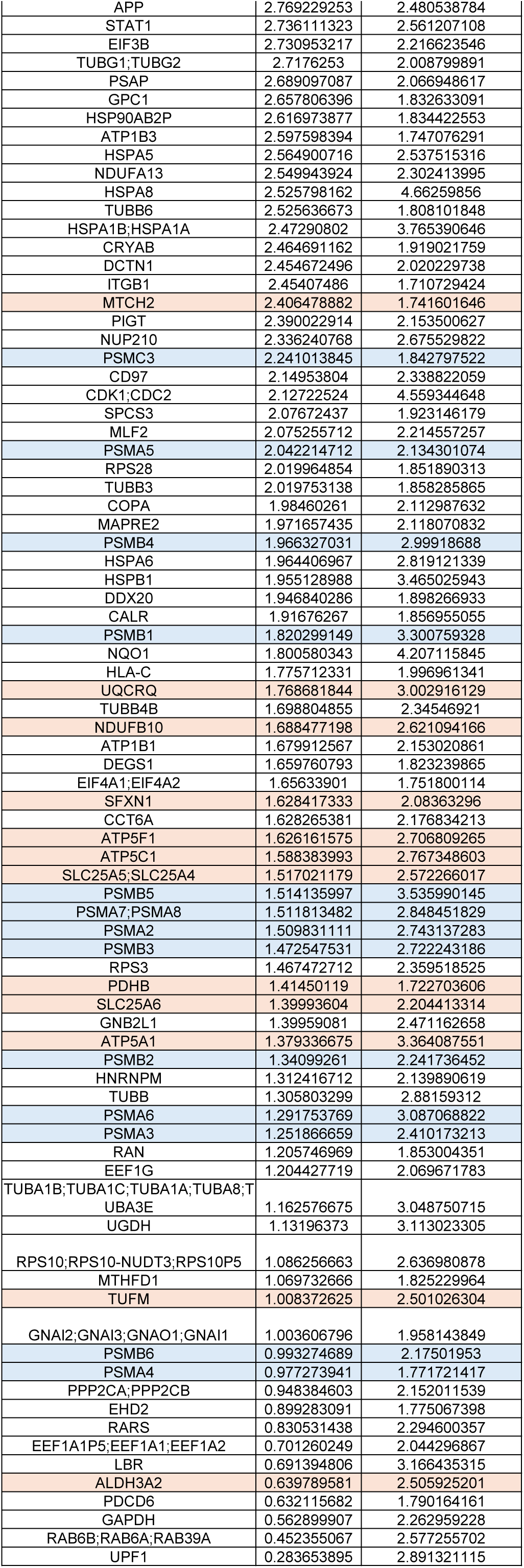
proteins copurifying with PLIN3B. List of protein significatively enriched in PLIN3B immunoprecipitation. Green = PLIN3B; red = mitochondrial proteins (MitoCarta3.0); blue = GOCC “proteasome complex” proteins.

The sequence of canonical PLIN3 was extensively matched in both samples with >70% sequence coverage and more than 40 peptide spectrum matches. Despite this, the QEQSYFMETV peptide was not identified. Upon manual inspection, no appreciable ion signal at the expected m/z and retention time was observed that would support the existence of this peptide.

A similar targeted experiment was performed on a total lysate of the same cells, digested in solution and without gel-based fractionation. No PLIN3B peptide was observed. PLIN3 sequence coverage was lower with in-solution digestion than after gel fractionation (data not shown).

### Public proteomics and ribosome profiling datasets provide no evidence for PLIN3B translation or protein detection

To assess evidence for PLIN3B protein expression, we analyzed publicly available datasets. Since PLIN3B is not present as an annotated protein in public proteomics databases, we queried the UniProt accession K7ERZ3, which contains the PLIN3B splicing junction. However, the junction-specific tryptic peptide QEQSYFMETV is not assigned a UniProt peptide accession and has never been observed in the Human PeptideAtlas 2025-01 build (https://peptideatlas.org/builds/human/), incorporating 806 analyzed proteomic datasets (Omenn et al., 2023).

To assess potential translation of *PLIN3B*, we compared ribosome profiling coverage of <u>ENST00000221957.9</u> (*PLIN3*) and <u>ENST00000589163.5</u> (*PLIN3B*/*K7ERZ3*) transcripts using Trips-Viz (Kiniry et al., 2021) (figure 6H). No ribosome-protected reads spanning the Chr19:4844794-4847816 junction, which represent the only PLIN3B-specific sequence, were detected.

Together, analyses of public proteomics and ribosome profiling databases provide no evidence of PLIN3B/K7ERZ3 protein detection or translation.

## Discussion

Because of our interest in investigating PLIN3 as a modulator of cellular energy metabolism, we serendipitously identified a transcript originating from a previously undescribed splicing event in *PLIN3* mRNA. Whereas PLIN3A primarily localizes to the cytosol and LDs, expression of PLIN3B resulted in mitochondrial localization.

Mitochondrial localization of PLIN3B was consistently observed across multiple detection strategies, including non-tagged PLIN3B detected with antibodies targeting both the 11-mers domain and the splicing junction, as well as N-terminal FLAG-tagged and internally ALFA-tagged PLIN3B. In contrast, localization of PLIN3B to LDs was less consistent and appeared to depend on experimental conditions and detection method. Notably, antibodies targeting the PLIN3 11-mers domain failed to detect PLIN3B and PLIN3A at LDs. The PLIN3 11-mers domain is naturally disordered and undergoes conformational changes upon membrane binding (Choi et al., 2023), which could reduce epitope accessibility and explain the apparent loss of antigenicity of LD-associated PLIN3 isoforms.

What determines PLIN3B mitochondrial localization remains unknown. Like PLIN3A, PLIN3B does not contain a cleavable mitochondrial targeting sequence, suggesting that its localization may rely on chaperone-mediated pathways. Chaperones such as HSP90 and HSPA8 have been implicated in mitochondrial protein targeting through TOM70-TOM40 (Young et al., 2003), and HSC70/HSPA8, together with its co-chaperone BAG2 (Qin et al., 2016). These were identified among PLIN3B interactors in our FLAG-based immunoprecipitation (table 3). Additionally, truncation analyses indicate that mitochondrial localization of PLIN3B depends on residues shared by both PLIN3 isoforms. Given that PLIN3A has previously been reported to interact with HSPA8 during lysosomal targeting and degradation (Kaushik and Cuervo, 2015), HSPA8 may be involved in distinct trafficking pathways of PLIN3 isoforms, and additional interactors may determine isoform-specific targeting. Importantly, PLIN3B mitochondrial localization was not associated with lysosomal or autophagic compartments, as assessed by LAMP2 colocalization, and ATG5 deficiency.

How alternative splicing contributes to these interactions remains unclear. AlphaFold predictions suggest a more open state of the PLIN3B 4HB, and the absence of the α/β domain that zips the 4HB (figure1F-K). In PLIN3A, the deletion of the α/β domain residues at the C-terminus of the protein was shown to modulate targeting by increasing LD association (Ajjaji et al., 2019). Phosphorylation of PLIN3A on Y251 has been reported to promote HSPA8 binding and lysosomal degradation (Yang and Rich, 2021). The Y251 residue lies within the 4HB domain affected by PLIN3B splicing (figure1F-K). It is therefore possible that structural alterations induced by the alternative splicing partially mimic phosphorylation-dependent conformational states, thus driving HSPA8 recruitment and mitochondrial import mediated by additional isoform-specific interactors. This interpretation is supported by correlative light–electron microscopy, which localizes PLIN3B signal within mitochondrial boundaries rather than at the MOM or organelle contact sites. Such structural changes could influence chaperone interactions (Choi et al., 2023) and engagement with the ubiquitin-proteasome complex (Fredrickson et al., 2011), thus contributing to the reduced stability of PLIN3B.

The reduced half-life of PLIN3B and its association with proteasomal subunits are consistent with low protein levels of overexpressed PLIN3B. Previous studies have described pathways linking unfolded or ubiquitinated proteins to the mitochondrial import machinery. Notably, the amyotrophic lateral sclerosis-associated protein TDP43 has been shown to be targeted to the mitochondrial import receptor TOMM70 through FUNDC1- and DNAJA2-dependent mechanisms (Ma et al., 2023). In parallel, FUNDC1 has been reported to interact with HSPA8/HSC70 to promote mitochondrial delivery of misfolded and ubiquitinated proteins, leading to their sequestration or degradation by mitochondrial proteases such as the LONP1 protease (Li et al., 2019). In this context, it is conceivable that PLIN3B mitochondrial localization could reflect engagement of protein quality control pathways. However, neither FUNDC1 nor LONP1 were identified among PLIN3B interactors (table 3), arguing against a direct involvement in these pathways. An alternative explanation is suggested by recent work from Oborská-Oplová et al. (Oborska-Oplova et al., 2025), who showed that constitutive mitochondrial targeting of yeast ribosomal subunits can be actively repressed by mitochondrial avoidance segments. In this framework, alternative splicing in PLIN3 exon 6 may generate structural rearrangements that functionally unmask residues capable of driving mitochondrial targeting, rather than creating a novel targeting signal per se. This interpretation supports the hypothesis that PLIN3 contains an intrinsic, but normally repressed, mitochondrial targeting potential that can be revealed under specific conditions regulating its secondary and tertiary structure. Our observations suggest the possibility of exploring how post-translational modifications and conformational changes within the PLIN3 α/β domain may regulate access to this latent mitochondrial targeting potential.

The absence of detectable endogenous PLIN3B protein remains a question in this study. Despite clear evidence for the *PLIN3B* splicing event and *PLIN3B* transcript expression, we did not obtain evidence for PLIN3B protein under basal conditions or following inhibition of proteasome- and lysosome-dependent degradation pathways. Notably, this absence of detectable PLIN3B protein persisted even when PLIN3B splicing was experimentally enhanced by antisense oligonucleotide treatment. One possible explanation is that low transcript abundance combined with the short half-life of PLIN3B results in protein levels below the detection limits of available approaches. However, the lack of PLIN3B detection in public proteomic datasets and ribosome profiling resources, in targeted proteomics analyses directed against the unique PLIN3B splicing junction peptide, and following perturbation of multiple protein degradation pathways argues against this possibility.

An alternative explanation is that *PLIN3B* translation is actively repressed. Because the exogenously expressed *PLIN3B* coding sequence is efficiently translated in transfected cells, regulatory elements outside the coding region, such as PLIN3B 3’UTR, may contribute to translation control. Notably, the *PLIN3* 3’UTR overlaps with the antisense transcript *PLIN3-AS1* (ENSG00000267484), raising the possibility that antisense-mediated mechanisms influence isoform-specific mRNA stability and translation. Further investigation of *PLIN3A* and *PLIN3B* mRNA stability and translation will be required to address these questions.

Finally, although less likely, the possibility that the *PLIN3B* cDNA represents a rare or artifactual amplification product cannot be completely excluded. More generally, this underscores the limitations of transcript annotation for inferring full-length protein-coding potential, particularly for low-abundance or partially characterized splice variants. One potential source of such an artefact would be chimeric amplification involving *PLIN3* and *K7ERZ3* transcripts, the latter of which remains incompletely annotated at its 5’ end. Alternative splicing of exon 6 could occur simultaneously with additional RNA processing events in *K7ERZ3*, resulting in a non-coding transcript (Herzel et al., 2018). Several observations argue against this explanation, including the detection of PLIN3B using multiple primer pairs and the amplification of the full-length cDNA following antisense oligonucleotide-mediated splicing modulation. Definitive exclusion of an amplification artefact, however, would require PCR-independent RNA detection approaches, such as Northern blotting, RNase protection assays, or direct RNA sequencing (Jain et al., 2022).

This study describes the identification and characterization of a previously unrecognized alternative splicing event in *PLIN3* that generates the *PLIN3B* transcript. Although this transcript is readily detectable and can be modulated by antisense oligonucleotides, we did not obtain evidence for endogenous PLIN3B protein expression under any of the experimental conditions tested. When expressed exogenously, however, the PLIN3B coding sequence encodes an unstable protein that localizes predominantly within mitochondria, in contrast to the LD-associated localization of PLIN3A. Together, these findings demonstrate that an alternative splicing event in *PLIN3* can generate a transcript whose coding sequence is sufficient to alter intracellular targeting and protein stability, yet does not give rise to a detectable endogenous protein product. The conservation of residues that mediate mitochondrial targeting between the two isoforms raises the question of how constitutive mitochondrial targeting may be masked in PLIN3. More broadly, this work provides a rigorous framework to distinguish transcript diversity from stable protein expression, and emphasizes the need for systematic experimental evaluation of the functional consequences of alternative splicing events.

### Contributions

Axel Aguettaz: conceptualization, data curation, formal analysis, investigation, methodology, project administration, validation, visualization, writing – original draft, review & editing. Yoan Arribat: conceptualization, methodology, writing – review & editing. Alexandra Schmitt, Aminata Gerihanov, Jocelyn Fleurimont, Vithusan Vijayatheva: investigation, validation. Jean Daraspe: methodology. Cassandra Tabasso: investigation, writing – original draft, review & editing. Sylviane Lagarrigue: conceptualization, writing – review & editing. Ernesto Picardi: methodology, investigation, writing – review & editing. Sarah Cohen: methodology, investigation, writing – review & editing. Christel Genoud: resources, supervision, writing – review & editing. Francesca Amati: conceptualization, formal analysis, funding acquisition, project administration, resources, supervision, validation, visualization, writing – original draft, writing – review & editing.

## Acknowledgments

We thank Manfredo Quadroni and the personnel of the Protein Analysis Facility, Danny Labes and Mariela Castelblanco Castelblanco at the Flow Cytometry Facility, as well as the Cell Imaging Facility of the Faculty of Biology and Medicine, University of Lausanne. We thank Thomas Langer of the Max Planck Institute for Biology of Ageing, Cologne, Germany, for sharing the HEK293 YME1L1 KO and control cell lines. We also thank Susmita Kaushik and Ana Maria Cuervo of the Einstein University, NY, for assistance with the studies in ATG5KO cells. We are also grateful to Allison Skinkle at the University of North Carolina at Chapel Hill, NC, for her support with multispectral imaging.

This study was supported by the Swiss National Science Foundation (grants: 320030_170062 and 310030_188789 to FA) and the La Fondation suisse de recherche sur les maladies musculaires (FSRMM, to FA).

## Data availability

Data are available in the manuscript and supplemental material. Proteomics data are openly available in ProteomeXchange at accession number PXD066308 and PXD066314. All public data used in this article are openly available. URLs accessible at time of submission are given in the methods section.

## Supplemental Figure legends

**Supplemental figure 1.**
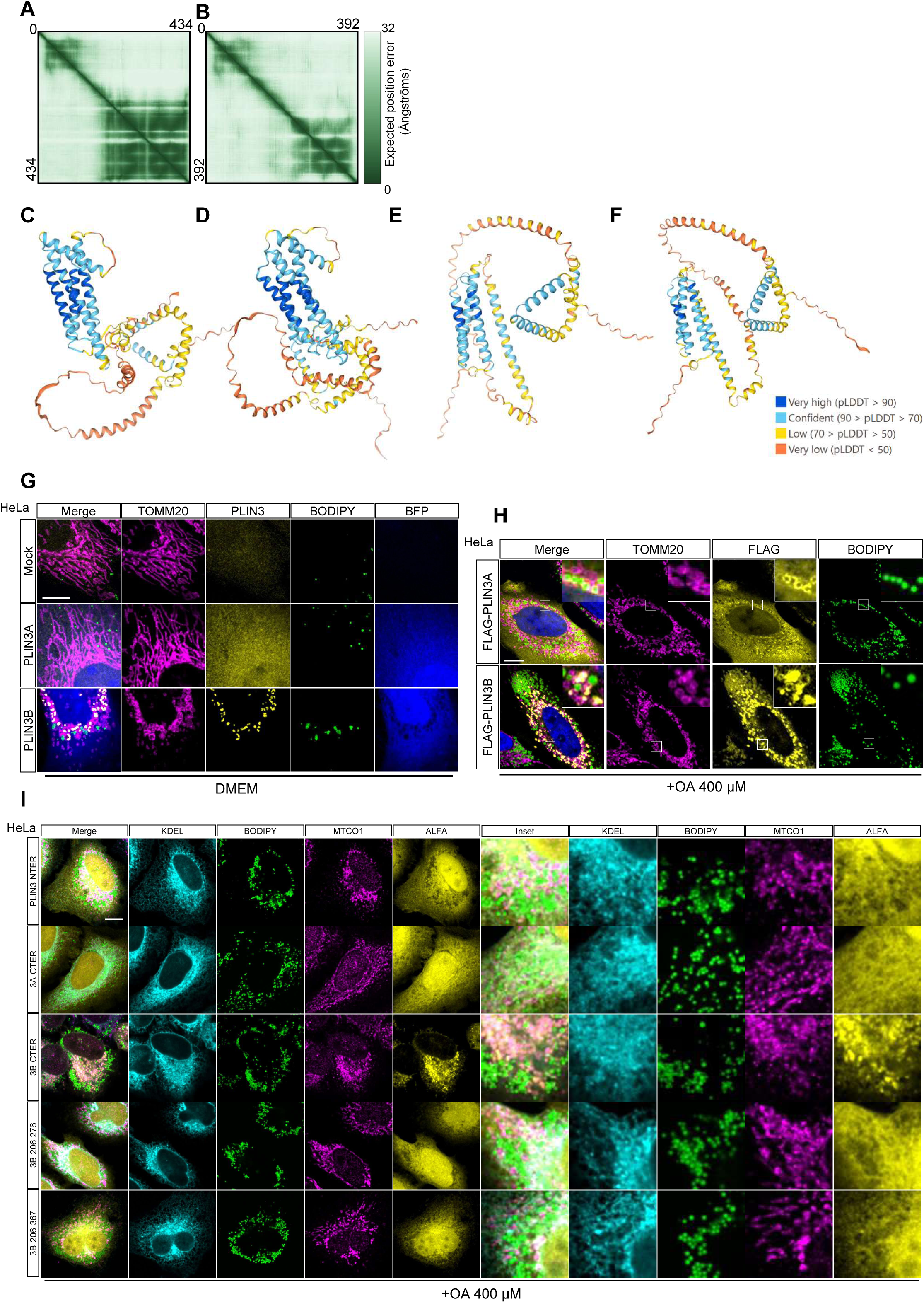
**A-B)** Predicted aligned error of PLIN3A **(A)** and PLIN3B **(B)** proteins from AlphaFold2 prediction. X and Y axis correspond to the position of individual residues in PLIN3A and PLIN3B chains. **C-F)** Predicted structure of PLIN3A **(C)**, PLIN3B **(D)**, PLIN3A-ALFA **(E)** and PLIN3B-ALFA **(F)**. Color code corresponds to predicted local distance difference test (pLDDT). **G)** Representative immunocytochemistry confocal images of HeLa cells transfected with PLIN3A-P2A-BFP or PLIN3B-P2A-BFP plasmids. Cells were stained with a PLIN3 antibody, with antibody against the mitochondrial marker TOMM20 and with BODIPY 493/503. Images were acquired with LSM780. Scale bar=10 µm. N=3 independent experiments. **H)** Representative immunocytochemistry confocal images of HeLa cells transfected with 3XFLAG-PLIN3A or 3XFLAG-PLIN3B plasmids and incubated with OA. Cells were stained with a PLIN3 antibody, with antibody againt the mitochondrial marker TOMM20 and with BODIPY 493/503. Images were acquired with LSM780. Scale bar=10 µm. N=3 independent experiments. **I**) Representative immunocytochemistry confocal images of HeLa cells transfected with different ALFA-tagged PLIN3A and PLIN3B fragments and incubated with OA. Cells were stained with an anti-ALFA antibody, with antibody against the mitochondrial marker MTCO1, with antibody against the ER marker KDEL and with BODIPY 493/503. Images were acquired with LSM780. Scale bar=10 µm. N=3 independent experiments.

**Supplemental figure 2.**
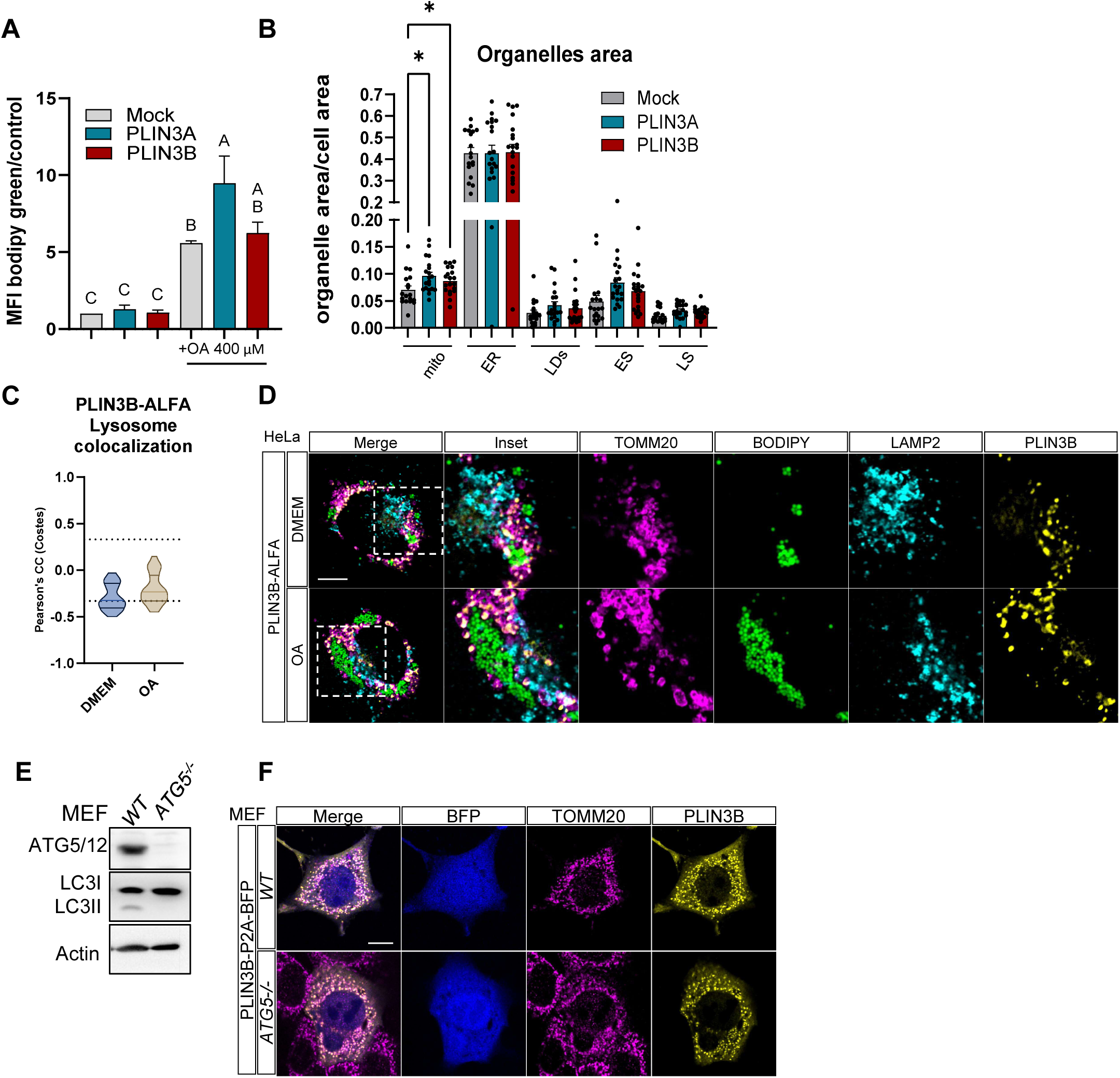
**A)** Flow cytometry analysis of HeLa cells transfected with PLIN3A-P2A-BFP and PLIN3B-P2A-BFP, with or without OA incubation. Cells were stained with BODIPY 493/503. Error bar=mean ± SEM. One-way ANOVA. Distinct letters (A, B, C) indicate significant differences between groups (significance was set at p<0.05). N=3 independent experiments. **B)** Quantification of relative organellar surface area in Mock, PLIN3A-P2A-BFP and PLIN3B-P2A-BFP-transfected cells in multispectral imaging experiment. Error bar=mean + SEM. Mixed-effects analysis, with the Geisser-Greenhouse correction, followed by two-stage linear step-up procedure of Benjamini, Krieger and Yekutieli with individual variances computed for each comparison. Mito=mitochondria, LS=lysosomes, ES=endosomes. N=18-20 cells per condition from 2 independent experiments **C)** Quantification of Pearson’s correlation coefficient of ALFA and LAMP2 cytosolic signal. Image thresholding was performed with Costes automatic threshold. n=28 cells per condition from 3 independent experiments. **D)** Representative immunocytochemistry confocal images of HeLa cells transfected with PLIN3B-ALFA and incubated in normal DMEM or in OA supplemented DMEM. Cells were stained with an anti-ALFA antibody, with antibody against the lysosomal marker LAMP2, and with BODIPY 493/503. Images were acquired with Stellaris 8CLSM. Scale bar=10 µm. N=3 independent experiments. **E)** Representative western blot from WT and *ATG5^-/-^* MEF. Expression of ATG5/12, LC3 and Actin was assayed. N=3 independent experiments. **F)** Representative immunocytochemistry confocal images of MEFs cells transfected with PLIN3B-P2A-BFP. Cells were stained with antibody against the mitochondrial marker TOMM20 and with anti-PLIN3 antibody. Scale bar=10 µm.

**Supplemental figure 3.**
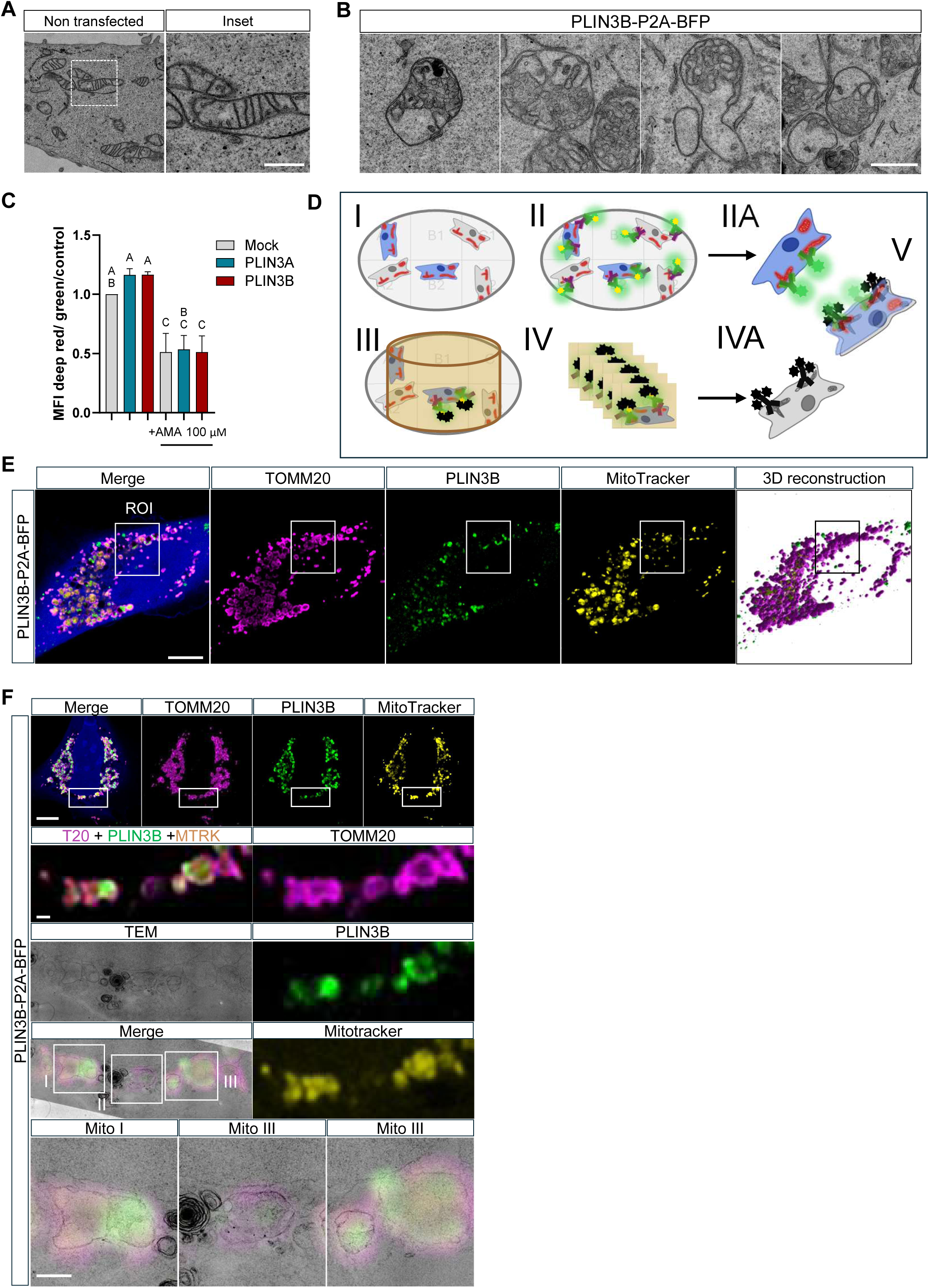
**A)** Representative electron micrographs of mitochondria from BFP negative HeLa cells. Scale bar=500 nm. **B)** Additional electron micrographs of mitochondria from cells transfected with PLIN3B-P2A-BFP. Scale bar=500 nm. **C)** Flow cytometry analysis of HeLa cells transfected with PLIN3A-P2A-BFP and PLIN3B-P2A-BFP, with or without the respiratory complex III inhibitor Antimycin A (AMA). Cells were stained with MitoTracker Deep Red FM and MitoTracker Green FM. Error bar=mean + SEM. One-way ANOVA. Distinct letters (A, B, C) indicate significant differences between groups (significance was set at p<0.05). N=3 independent experiments. **D)** Schematics of FNG-CLEM protocol: **I)** Cells are seeded on Mattek dishes, transfected and stained with MitoTracker Red CMXRos. **II)** Cells are fixed and permeabilized with LN2. Immunostaining is performed with TOMM20 and PLIN3B antibodies. A FluoroNanogold secondary antibody is used for the detection of PLIN3B 4D72 antibody. **IIA)** Cells are imaged with CLSM. 3D stacks (200nm z-step) are acquired for the whole cell volume. **V)** Sample is processed, nanogold is enhanced with colloidal silver and embedded in resin. **E)** Serial ultrathin sections (100nm) are obtained for ROIs corresponding to the desired cells. **IVA)** Acquisition of electron micrographs. **V)** 3D reconstruction and CLSM images are rotated and flipped and correlated with electron micrographs. **E)** Representative immunocytochemistry image and 3D reconstruction of PLIN3B-P2A-BFP transfected HeLa cell in FNG-CLEM protocol. Staining was performed with MitoTracker Red CMXRos, PLIN3B 4D72 antibody and with anti-TOMM20 antibody. FNG conjugated secondary antibody was used to target PLIN3B. ROI corresponding to the area analysed in the **figure 3E-G**. Scale bar=10 µm. **F)** Representative immunocytochemistry image and TEM section of FNG-CLEM protocol, without silver enhancement of gold particles. Staining was performed with MitoTracker Red CMXRos, PLIN3B 4D72 antibody and with anti-TOMM20 antibody. FNG conjugated secondary antibody was used to target PLIN3B. Scales bars, from top to bottom= 10, 1, 1 µm.

**Supplemental figure 4.**
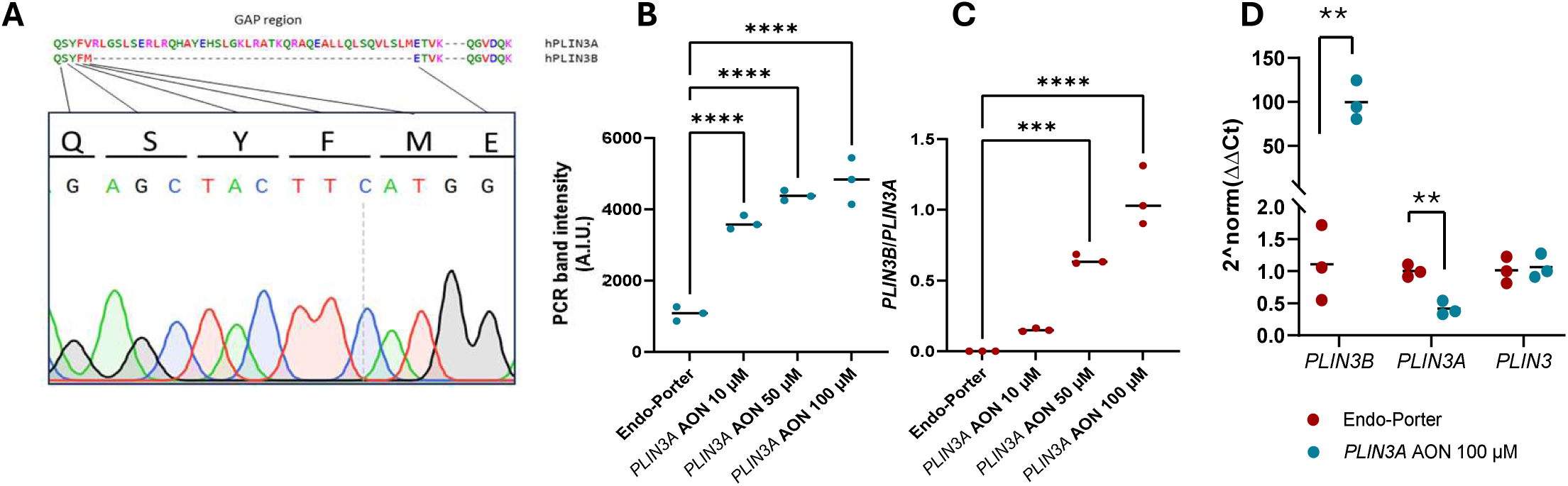
**A)** Representative image of Sanger sequencing on *PLIN3B* junction amplicon from *PLIN3A* AON treated HUVEC cells in **figure 5D**. **B-C)** Quantification of *PLIN3B* junction amplicon band intensity **(B)** and *PLIN3B/PLIN3A* ORF amplicon band intensity ratio **(C)** from **figure 5A**. ∗∗∗∗p < 0.0001, ∗∗∗p < 0.001, ∗∗p < 0.01, ∗p < 0.05 (one-way ANOVA), N=3 independent experiments. **D)** QPCR of *PLIN3B*, *PLIN3* and *PLIN3A* levels in AON treated HeLa cells. ∗∗∗∗p < 0.0001, ∗∗∗p < 0.001, ∗∗p < 0.01, ∗p < 0.05 (unpaired t-test). N=3 independent experiments.

**Supplemental figure 5.**
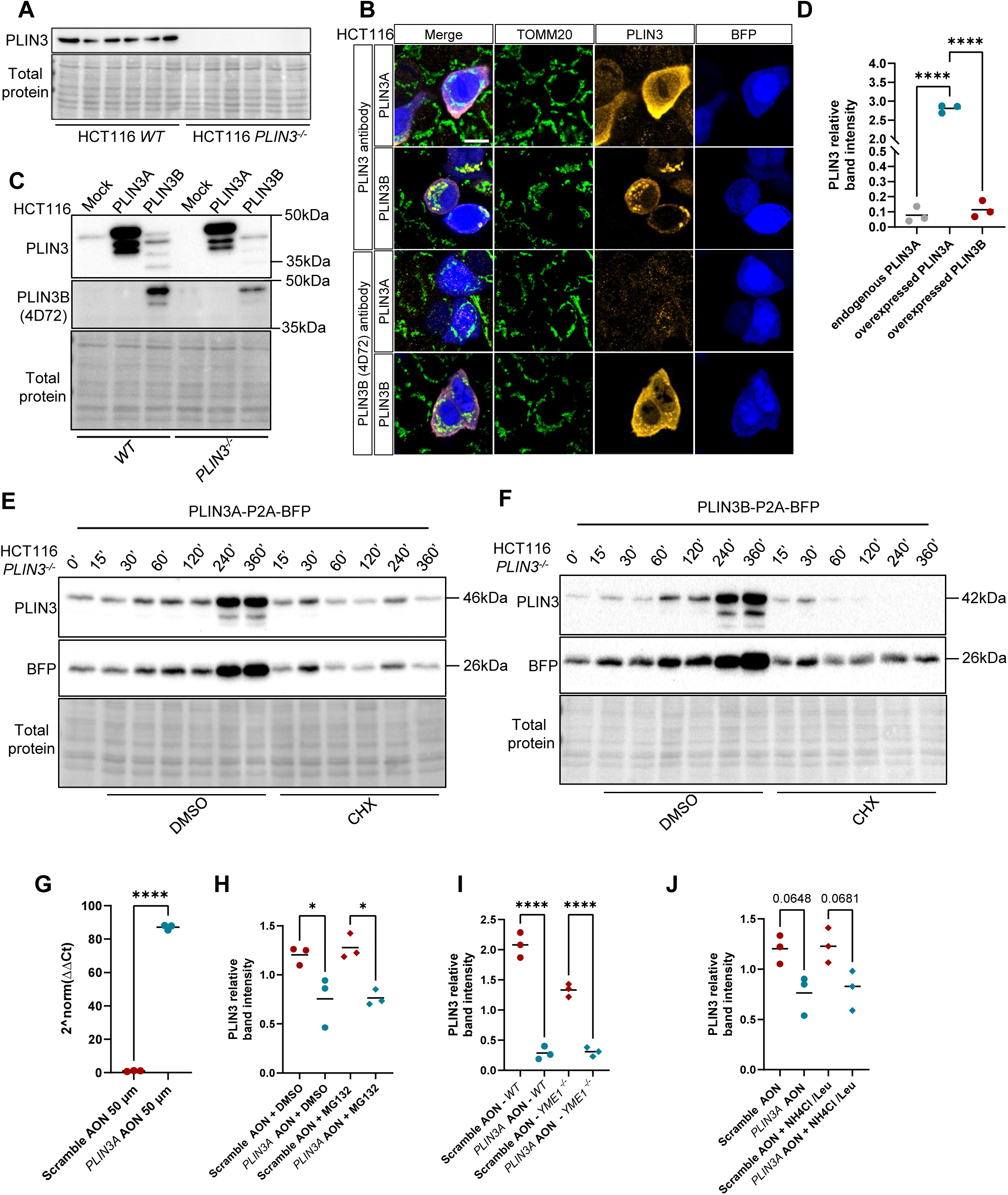
**A)** Representative western blot of *WT* and CRISPR-Cas9 generated *PLIN3^-/-^ KO* HCT116 cells. Expression of PLIN3 was assayed. N=6 *KO* lines tested. **B)** Representative immunocytochemistry of HCT116 cells transfected with PLIN3A-P2A-BFP and PLIN3B-P2A-BFP. Cells were stained with antibody targeting TOMM20 mitochondrial marker, with PLIN3 antibody and with the PLIN3B-specific 4D72 antibody from hybridoma supernatant. N=3 independent experiments. Scale bar=10 µm. N=3 independent experiments. **C)** Representative western blot of HCT116 cells transfected with PLIN3A-P2A-BFP and PLIN3B-P2A-BFP. Expression of PLIN3 was assayed with the isoform non-specific antibody and with the PLIN3B-specific 4D72 antibody from hybridoma supernatant. N=3 independent experiments. **D)** Quantification of PLIN3A and PLIN3B levels in **supplemental figure 5C**. ∗∗∗∗p < 0.0001, ∗∗∗p < 0.001, ∗∗p < 0.01, ∗p < 0.05 (Ordinary one-way ANOVA), N=3 independent experiments. **E-F)** Representative western blot of CHX pulse-chase in PLIN3A-P2A-BFP **(D)** and PLIN3B-P2A-BFP **(E)** transfected HCT116 *PLIN3^-/-^* KO cells. Expression of PLIN3 and BFP was assayed. N=3 independent experiments. **G)** qPCR of *PLIN3B* levels in HCT116 cells treated with *PLIN3A* AON. Error bar=mean, ∗∗∗∗∗p < 0.0001, ∗∗∗p < 0.001, ∗∗p < 0.01, ∗p < 0.05 (unpaired *t* test), N=3 independent experiments. **H-J)** Quantification of PLIN3 band intensity in figure 6D **(H**), 6E **(I)** and 6F **(J)**. Error bar=mean. ∗∗∗∗p < 0.0001, ∗∗∗p < 0.001, ∗∗p < 0.01, ∗p < 0.05 (Ordinary one-way ANOVA), N=3 independent experiments.

## Material and methods

### Cell culture and handling

Cell lines were obtained from the sources listed in **table 4**. HUVEC cells were cultured with EGM2 medium (Lonza, Basel, Switzerland, CC-3162) in 5%CO_2_, 5%O_2_ with bovine skin gelatin (Sigma-Aldrich, Saint-Louis, USA, G9382) coating. HeLa, HEK293 and HCT116 were cultured with DMEM (DMEM GlutaMAX (GIBCO, Waltham, USA, 31966–021) supplemented with 10% fetal bovine serum (FBS) (10270–016, GIBCO) and 100 IU/ml penicillin (15140122, GIBCO), 100 μg/ml streptomycin (15140122, GIBCO) in 5%CO_2_, 21%O_2_. Cells were trypsinized at subconfluence with Trypsin 0.05%, EDTA (Life technologies, Saint-Aubin, France, 25300054).

**Table 4:** cell lines used in this manuscript.

| Cell line | origin |
| --- | --- |
| HeLa | Sigma-Aldrich (93021013) |
| HEK293 *YME1L*<sup>-/-</sup> and control | Kind gift of Prof. Thomas Langer, Max Planck Institute for Biology of Ageing, Cologne, DE |
| HUVEC | Kind gift of Dr. Florent Allagnat, Department of Vascular Surgery, CHUV, Lausanne, VD, CH |
| HCT116 | Kind gift of Prof. Lluis Fajas Coll, Center for Integrative Genomics, University of Lausanne, VD, CH |
| MEF WT | Noboru Mizushima, Department of Biochemistry and Molecular Biology, University of Tokyo, JP |
| MEF ATG5 KO | Noboru Mizushima, Department of Biochemistry and Molecular Biology, University of Tokyo, JP |

For electron microscopy experiments, HeLa cells were cultured on 35 mm Dish with No. 1.5 Gridded Coverslip, 14 mm Glass Diameter (MatTek, Ashland, USA, P35G-1.5-14-C-GRD). For immunocytochemistry experiment, cells were cultured on 8 Well #1.5 polymer bottom µ-Slides with Ibitreat (Ibidi, Gräfelfing, Germany, 80806) or on No. 1 VWR® 12 mm Round Micro Cover Glasses, (VWR, Radnor, USA, 631-1577). Transfection was performed with Lipofectamine™ 3000 Transfection Reagent (Invitrogen, Waltham, USA, L3000015) for HeLa cells and with X-tremeGENE™ 360 Transfection Reagent (Roche, Basel, Switzerland, XTG360-RO) for HCT116 cell. For oleic acid treatment, 16 mg/ml of Fraction V fatty acid-free bovine serum albumin (Sigma-Aldrich, 10775835001) was diluted in complemented DMEM. Oleic acid (Sigma-Aldrich, O1383) was then solubilized in BSA-DMEM to a final concentration of 400 µM. MG132 (Sigma-Aldrich, 474787), Leupeptin (Sigma-Aldrich, L9783) and Antimycin A (Sigma-Aldrich, A8674) were resuspended in Dimethyl sulfoxide (Sigma-Aldrich, D8418). Ammonium Chloride (Sigma-Aldrich, A9434) was resuspended in double-distilled water. MitoTracker Green FM (Life technologies, M7514), MitoTracker Red CMXRos (Life technologies, M7512) and MitoTracker Deep Red FM (Life technologies, M22426) were resuspended in DMSO according to manufacturer’s protocol.

### Morpholino antisense oligonucleotide treatment

Cells were seeded in 6 well-plates (HUVEC, HeLa) or 12 well-plates (HCT116, HEK293) at 60% confluence. 24 h later, cells were treated with *PLIN3A* morpholino antisense oligonucleotide (Gene Tools, Philomath, USA, **table 2**), random Scramble morpholino 25-mers (Gene Tools), and 6 µM Endo-Porter (Gene Tools, 2922498000) according to manufacturer’s protocol and incubated 72 h. For proteomic analysis, cells were trypsinized and the pellet washed 3 times in PBS at 600 x g, snap frozen and stored at -70°C. For western blot (WB) and polymerase chain reaction (PCR) applications, samples were processed as described in the **RNA extraction and retrotranscription** and **cell lysis and protein quantification** sections.

### RNA extraction and retrotranscription

RNA extraction was performed with TRI Reagent® (Sigma-Aldrich, T9424) according to manufacturer’s protocol. In brief, cells were incubated with TRI Reagent® for 10’, followed by addition of Chloroform (Sigma-Aldrich, C2432), 10’ incubation and centrifugation at 16000 x g for 10’ at 4°C. The liquid phase was mixed with Isopropanol (Sigma-Aldrich, 34863) and incubated overnight (ON) at -20°C. 1 µl of Glycogen was added to the solution to visualize the RNA pellet, followed by centrifugation at 12000 x g for 10’ at 4°C. The pellet was washed 3 times in 75% ethanol (Carlo Erba Reagents, 4146072) in nuclease-free water (Thermo Fisher Scientific, Waltham, USA), air dried, then resuspended in 20 µl nuclease-free water and incubated at 55°C for 10’ to allow complete resuspension. The solution was quantified with NanoDrop™ One Microvolume UV-Vis Spectrophotometer (Thermo Fisher scientific). Retrotranscription was performed with GoScript Reverse Transcription Mix, Random Primers according to manufacturer’s protocol (Promega, Madison, USA, 2801). The Invitrogen™ Ambion™ FirstChoice™ Human Reference RNA panel (Invitrogen, AM6050) used for the PCR experiments was retrotranscribed with the same protocol.

### Endpoint PCR

PCR was performed with GoTAQ long PCR master mix (Promega, M4021) according to manufacturer’s protocol. For PLIN3B differential splicing domain primer pair (**table 2**), the master mix was complemented with 5 µM MgCl (Promega, AB351B). The PCR product was mixed with 6X Loading Dye (Thermo Scientific, R0611) and loaded on 2% agarose gel (Roche, 11388983001) in TBE with SYBR™ Safe DNA Gel Stain (Life technologies, S33102) at 1:5000 dilution. Gel electrophoresis was performed at 100 V for 30’. Gels were imaged with a BioDoc-it imaging system (Thermo Fisher scientific) and pictures were analyzed with ImageJ (Schneider et al., 2012). For amplicon sequencing, bands were purified with MinElute Gel Extraction kit (28604, Qiagen, Hilden, Germany) and sequenced by Sanger sequencing by Fasteris (Plan-Les-Ouates, Switzerland). Alignement was performed with Snapgene (San Diego, United States).

### QPCR

QPCR was performed with Power SYBR™ Green PCR Master Mix (Thermo fisher, 4367659) according to manufacturer’s protocol with a QuantStudio™ 5 Real-Time PCR machine (Thermo Fisher Scientific). Quantification was performed with the 2delta-deltaCt quantification technique (Livak & Schmittgen, 2001).

### Immunofluorescence and imaging

Cells were fixed in 4% formaldehyde (FA) (Sigma-Aldrich, 252549) in PBS (Dr. G. Bichsel AG, Interlaken, Switzerland) at room temperature (RT) for 20’, followed by permeabilization with 0.25% TritonX-100 (Sigma-Aldrich, X100) in PBS 1X solution for 10’. Blocking was performed with 2% bovine serum albumin (Sigma-Aldrich, A7030) in PBS 1X for 10’. Cells were then incubated 45’ with primary antibodies (**table 1**) in BSA 2% PBS solution or with hybridoma supernatants, washed thrice and then incubated with secondary antibodies (**table 1**) in BSA 2% PBS solution for 45’. When PLIN3B 4D72 antibody was used, primary antibodies were directly diluted in the hybridoma supernatant, and cells were incubated 45’ with the solution. Hoechst 33342 (Invitrogen, Eugene, Oregon, 62249) staining was performed at 1:8000 dilution in PBS for 5’ after secondary antibody incubation, followed by 3 washes with PBS. BODIPY™ 493/503 (Invitrogen, D3922) staining was performed at 1:1000 dilution in BSA 2% PBS solution for 10’, followed by 1 wash in PBS. Coverslip-seeded cells were mounted with Mowiol® 4-88 (Sigma-Aldrich, 81381) on SuperFrost Plus™ Adhesion Microscope Slides (Epredia, Basel, Switzerland, J7800AMNZ). Multiwell-seeded cells were mounted with 25% glycerol (Sigma-aldrich, G5516) in PBS. Images were acquired with LSM780 confocal laser-scanning microscope (Carl Zeiss, Oberkochen, Germany) with Plan Apochromat 63x /1.40 numerical aperture (NA) / oil immersion objective, with Leica Stellaris 8 CLSM (Leica Mikrosysteme GmbH, Vienna, Austria) with HC PL APO 63x/1.40 oil CS2 objective or with Leica TCS SP8 confocal microscope using HC PLAPO CS2 63x/1.40 oil objective.

Super resolution images were acquired with Elyra7 (Carl Zeiss) with Plan Apochromat 63x / 1.4 NA / oil / WD 0.19 mm objective in 3D Lattice illumination mode and deconvolved using Zen 3D SIM^2^ processing (Carl Zeiss). All oil immersion microscopy was performed with Immersol 518F (Carl Zeiss).

### Confocal microscopy colocalization analysis

.lsm and .lif images obtained from LSM780 or Stellaris 8 were opened with ImageJ. Cytosolic fluorescence signal was isolated from nuclear signal by creating a mask. Images were denoised using the “Smooth” function and an intensity value of 5 was removed from all channels using the “subtract” function. Colocalization was analysed with the Coloc2 plugin (https://imagej.net/plugins/coloc-2) using Costes automatic thresholding.

### Cell Lysis and protein quantification

Cells were lysated with 1% NP40 IGEPAL® CA-630, NP40 equivalent (Sigma-Aldrich I8896), 150 mM Sodium chloride (Thermo-Fischer Scientific, S/3120/60), 5 mM EDTA (Sigma-Aldrich, ED), 50 mM Tris (Sigma-Aldrich, T1503) Lysis buffer in PBS. The lysis buffer was complemented with 1 mM Sodium orthovanadate (Sigma-Aldrich, S6508), 1 mM β-glycerophosphate (Sigma-Aldrich, 50020), 50 mM Sodium Fluoride (Sigma-Aldrich, 201154), 2.5 mM Sodium pyrophosphate (Sigma-Aldrich, P8010) and 1x Protease inhibitor cocktail (Sigma-Aldrich, P8340). In brief, cells were washed 3 times in PBS and incubated on ice 5’ with complemented lysis buffer, scraped and incubated 1 h at 4°C under rotation. Samples were centrifuged at 16000 x g for 20’ at 4°C. Proteins in supernatants were then quantified with BCA Protein Assay Kit (Sigma-Aldrich, 71285-M). Protein solution was diluted in Laemmli buffer: Tris 62.5 mM, 2% sodium dodecyl sulfate (SDS) (Sigma-Aldrich, L3771), 10% glycerol, 1% 2-mercapto-ethanol (Acros Organics, Geel, Belgium, 125472500), Bromophenol Blue (Thermo fisher, BP115-25).

### Western blot

5-20 μg of proteins from each lysate were loaded in 12% SDS-Page gels. Transfer was performed on methanol-activated PVDF membrane (Bio-Rad Laboratories, Hercules, USA) in tris-glycine (Biosolve Chimie, Dieuze, France) 20% ethanol buffer. Membranes were stained with Pierce MemCode™ Reversible Protein Stain Kits for WB (Thermo Scientific, 24585). After blocking 5% w/v skim milk powder (Sigma-Aldrich, 70166), 0.1% Tween20 (Thermo Fisher scientific, T10466) in PBS, primary antibodies (**table 1**) were incubated ON at 4°C. HRP-linked secondary antibodies (**table 1**) were diluted in 5% w/v skim milk powder, 0.1% Tween20 in PBS and incubated for 2 h at RT. Signal acquisition was performed on Biorad Chemidoc (ChemiDoc XRS+, Bio-Rad Laboratories) using WesternBright (Advansta, San Jose, USA, K-12045) or WesternBright Sirius (Advansta, K-12043) and quantified by densitometry using ImageJ. Protein levels werenormalized to the total protein concentration corresponding to MemCode™ staining. In **figure 2B**, band intensity was normalized to the sum of cytosolic and mitochondrial bands.

For **supplemental figure 2E**, cell lysates were prepared in RIPA buffer (150mM sodium chloride, 1% NP-40, 0.5% sodium deoxycholate, 0.1% SDS, 50mM Tris pH8) containing protease inhibitors (100μM AEBSF, 100μM leupeptin, 10μM pepstatin, 1mM EDTA). Protein concentration was measured by the Lowry method and samples subjected to SDS-PAGE and immunoblot using standard procedures. Proteins were visualized by using peroxidase-conjugated secondary antibodies and Western Lightning Plus (PerkinElmer, Shelton, USA) in a G-BOX Chemi XX6 (Imgen, Alexandria, USA).

### Gateway® cloning and plasmids

For the plasmids PLIN3A-P2A-BFP, PLIN3B-P2A -BFP and 3xFLAG-PLIN3B, coding sequences were cloned from human muscle cDNA and amplified by PCR with ATTB flanking sequences for the Gateway cloning system (table 2). PCR products were purified on agarose gels (MinElute Gel Extraction kit, Qiagen, Hilden, Germany, 28604) and inserted in Gateways pDon221 vector by recombination using BP clonase II enzyme mix (Invitrogen, 11789). CDS were transferred in destination vectors pcDNA-pCI-CMV-3xFLAGN (Grepper et al., 2024) or pCS Dest2 Attb1-3 (Addgene, Watertown, USA, 22424) + pDonR2R3-p2a-BFP with LR clonase II enzyme mix (Invitrogen, 11791). ALFA-tagged PLIN3A and PLIN3B plasmids were generated by Vectorbuilder (Neu-Isenburg, Germany) and are based on the sequences cloned from human muscle samples and on the ALFA-tag sequence (Nanotag biotechnologies, Göttingen, Germany).

### PLIN3 Knock Out line generation

PLIN3 Knock Out (KO) was performed with Crispr-Cas9 techniques using TIP47 CRISPR/Cas9 KO Plasmid (h) (Santa Cruz BTC, sc-405514) following Manufacturer’s protocol. Selection was performed with 2 µg/ml doxicyclin (Sigma-Aldrich, D9891) for 10 days. Single cells were isolated and screened with WB for PLIN3 presence. KO clones were transfected with TIP47 HDR Plasmid (h) (Santa Cruz Biotechnologies, sc-405514-HDR) and recombinant cells were selected for further experiments.

### PLIN3B antibody generation

Antibody production was performed by Genscript (Piscataway, USA). In brief, BALB/c Mice were immunized with KLH-conjugated QEQSYFMETVKQGVDC (Genscript) PLIN3B junction peptide. Spleen B-cells were fused with SP2/0 Myeloma cells. Hybridoma cells supernatants were screened with ELISA against the immunization peptide and PLIN3 fragments corresponding to 5’ and 3’ of the splicing junctions. Supernatants from clones recognizing exclusively PLIN3B junction peptide were tested *via* WB against cell lysates expressing PLIN3A-P2A-BFP or PLIN3B-P2A-BFP.

### Cycloheximide pulse chase

*PLIN3^-/-^* KO HCT116 cells were transfected with PLIN3A-P2A-BFP or PLIN3B-P2A-BFP plasmids. 24 h post-transfection medium was replaced with fresh DMEM containing DMSO (Sigma-Aldrich, D8418) or Cycloheximide (Sigma-Aldrich, C7698) at 100 µg/ml concentration. At the indicated timepoints, samples were processed as in **cell lysis**, **protein quantification** and **WB** sections.

### Multispectral analysis

U2OS cells were transfected with pCMV-PLIN3A-p2a-BFP or pCMV-PLIN3B-p2a-BFP to select PLIN3-expressing cells by indirect visualization of BFP. U2OS cells were cultured in 8-well chambered coverglass (Cellvis) coated with 10 μg/ml fibronectin (Millipore). Cells were prepared for multispectral imaging as previously described. Briefly, cells were transfected to express PLIN3A-p2a-BFP or PLIN3B-p2a-BFP, LAMP1-CFP (late endosomes/lysosomes), mito-EGFP (mitochondria), and mApple-Rab5a-7 (early endosomes; Addgene #54944, from Michael Davidson), and then incubated with Bodipy 665/676 (50 ng/ml, Life Technologies) for 16 h to label LDs. Images were acquired on a Zeiss 880 laser confocal scanning microscope equipped with a 32-channel multi-anode spectral detector (Carl Zeiss) using a 63×/1.4 NA objective lens at 37 °C and 5% CO₂. All fluorophores were excited simultaneously using 458, 514, 561, and 640 nm lasers, and images were collected onto a linear array of 32 photomultiplier tube elements in lambda mode at 9.7 nm bins. The emission spectra of the fluorophores were defined using images from singly labeled cells, and images from multiply labelled cells were subjected to linear unmixing using Zen software (Carl Zeiss). For image analysis, a specialized masking protocol was designed in the open-source software CellProfiler to detect morphologically distinct organelles with high specificity (https://cellprofiler.org). Detection was individually optimized for each organelle by tuning the level of Gaussian blurring, carefully adjusting thresholding parameters, and applying pixel-based size cutoffs. Masks were then generated for cells and their detected organelles. Organelle number and size were calculated from these masks using a custom MATLAB pipeline (AnalyzeMultispectral, available at: https://github.com/TimXQi/Cohen-Lab). Organelle area fraction was calculated by dividing the number of pixels in the organelle mask by the number of pixels in the cell mask. Contact area fraction was calculated by generating an overlap image between two distinct organelle masks, summing the number of overlapping pixels, and dividing by the number of pixels in the first organelle mask. All reported values are medians.

### Cell fractionation

HeLa cells were trypsinized and centrifuged at 300 x g for 5’ at 4°C. The pellet was washed 2 times in PBS, resuspended in IB buffer (Jha et al., 2016) complemented with proteases and phosphatases inhibitors (1mM Na3VO4, 1mM b-glycerophosphate, 50 mM NaF, 2.5 mM Sodium pyrophosphate and 1x Proteases inhibitor cocktail (Sigma-Aldrich, P8340)) and homogenized with 30 strokes at 2500 rpm with a teflon–glass dounce homogenizer. All steps were performed at 4°C. After 3 subsequent centrifugation rounds at 1000 x g for 5’, the supernatant was collected and centrifuged at 10’000 x g for 10’. Supernatant (cytosolic fraction) and pellet (mitochondria fraction) were separated. The mitochondria fraction was resuspended in IB buffer, centrifuged again at 10’000 x g, resuspended and quantified with BCA Protein Assay Kit (Sigma-Aldrich71285). The cytosolic fraction was centrifuged at 10’000 x g, and the supernatant was separated from residual mitochondria.

### Co-Immunoprecipitation

Cleared protein extract (see **cell lysis and protein quantification** section) of Mock transfected or 3XFLAG-PLIN3B transfected HeLa cells were immunoprecipitated with Dynabeads™ Protein A magnetic beads (Invitrogen, 10002D) according to the manufacturer’s protocol. In brief, 1 mg of proteins were incubated with M2 FLAG antibody (Sigma-Aldrich, F1804-M2)-conjugated beads for 4 h at 4°C under rotation. Beads were washed 3 times in 1:1 1% IGEPAL CA-630 lysis buffer:PBS solution and 3 times in ice-cold PBS. Proteins were eluted with pH-based elution with a 2.5% Ammonia (Sigma-Aldrich, 5438300100) solution in 3 consecutive 10’ elutions at 4°C. Pooled elutions were stored at -70°C prior to proteomic analysis.

### Sample preparation and proteomic analysis of PLIN3B co-immunoprecipitations

Elutes from FLAG immunoprecipitation were lyophilized to dryness. Unless otherwise specified, all chemicals for proteomics steps were from Sigma-Aldrich. Protein digestion followed a modified version of the iST method (Kulak et al., 2014) (miST). Samples were redissolved in 50 μl miST lysis buffer (1% Sodium deoxycholate, 100 mM Tris pH 8.6, 10 mM DTT), heated 5’ at 95°C and diluted 1:1 (v:v) with water. Reduced disulfides were alkylated by adding ¼ vol. of 160 mM chloroacetamide (32 mM final) and incubating for 45’ at RT in the dark. Samples were adjusted to 3 mM EDTA and digested with 0.5μg Trypsin/LysC mix (Promega V5073) for 1 h at 37°C, followed by a second 1 h digestion with an additional 0.5μg of proteases. To remove sodium deoxycholate, two sample volumes of isopropanol containing 1% trifluoroacetic acid (TFA) were added to the digests, and the samples were desalted on a strong cation exchange (SCX) plate (Oasis MCX micro-elution plate; Waters Corp., Milford, USA, 186001830B) by centrifugation. After washing with isopropanol/1%TFA, peptides were eluted in 200 μl of 80% MeCN, 19% water, 1% (v/v) ammonia, and dried by centrifugal evaporation.

Tryptic peptide mixtures were resuspended in 25 μl of 0.1% formic acid and 5 μl were injected on a Vanquish Neo nanoHPLC system interfaced via a nanospray Flex source to a high resolution Orbitrap Exploris 480 mass spectrometer (Thermo Fisher). Peptides were loaded onto a trapping microcolumn PepMap100 C18 (5 mm x 1.0 mm ID, 5 μm, Thermo Fisher, 160434) before separation on a C18 custom packed column (75 μm ID × 45 cm, 1.8 μm particles, Reprosil Pur, Dr. Maisch, Ammerbuch, Germany), using a gradient from 2 to 80% acetonitrile in 0.1% formic acid for peptide separation at a flow rate of 250 nl/min (total time: 130’). Full MS survey scans were performed at 120’000 resolution. A data-dependent acquisition method controlled by Xcalibur software (Thermo Fisher Scientific) was used that optimized the number of precursors selected (“top speed”) of charge 2^+^ to 5^+^ while maintaining a fixed scan cycle of 2s. Peptides were fragmented by higher energy collision dissociation (HCD) with a normalized energy of 30% at 15’000 resolution. The window for precursor isolation was of 1.6 m/z units around the precursor and selected fragments were excluded for 60s from further analysis.

Data files were analysed with MaxQuant 1.6.14.0 (Cox & Mann, 2008a) incorporating the Andromeda search engine (Cox & Mann, 2008b). Cysteine carbamidomethylation was selected as fixed modification while methionine oxidation and protein N-terminal acetylation were specified as variable modifications. The sequence databases used for searching were the human reference proteome (RefProt_Homo_sapiens_20210620.fasta) based on the UniProt database, June 2021 version (Bateman et al., 2021), and a “contaminant” database containing the most usual environmental contaminants and enzymes used for digestion (keratins, trypsin, etc). Mass tolerance was 4.5 ppm on precursors (after recalibration) and 20 ppm on MS/MS fragments. Both peptide and protein identifications were filtered at 1% FDR relative to hits against a decoy database built by reversing protein sequences. The “match between runs” function of MaxQuant was activated. LFQ values (Cox et al., 2014) were used for all subsequent calculations.

All subsequent analyses were done with the Perseus software package (version 1.6.15.0) (Tyanova et al., 2016). Contaminant proteins were removed and LFQ values were log2-transformed. After assignment to groups, only proteins quantified in at least 3 samples of one group were kept. After missing values imputation (based on normal distribution using Perseus default parameters), t-tests were carried out among conditions, with permutation-based FDR correction for multiple testing (adjusted p-value threshold <0.05). Imputed values were later removed. GOCC analysis in **figure 4** was performed with ShinyGO0.77 (Xijin Ge et al., n.d.). Significantly enriched proteins in FLAG-PLIN3B were analyzed using all proteins detected in the analysis as background, pathway size between 10 and 1000 terms, FDR-filtered (q-val<0.05). Heatmaps were generated with Morpheus (RRID:SCR_017386, Broad Institute).

### GTEx database analysis

To evaluate the frequency of the splice event from different human healthy tissues, GTEx RNAseq experiments were downloaded from the database of Genotypes and Phenotypes (dbGaP, https://dbgap.ncbi.nlm.nih.gov/home/) (Mailman et al., 2007) under the accession number phs000424.v7.p2 in SRA format and converted in standard fastq format using the fastq-ump from SRA toolkit (https://trace.ncbi.nlm.nih.gov/Traces/sra/sra.cgi?view=software). RNAseq reads were aligned to the human genome (hg19/GRCh37) using STAR version 2.5 (Dobin et al., 2013) providing known gene annotations from Gencode version 31 (Frankish et al., 2019). Alignments were converted in BAM format by SAMtools72. The number of reads supporting Junction 3 (Chr19:4844794-4847690, generating ENSE00000664968 exon, *PLIN3A*) and Junction 4 (Chr19:4844794-4847816, generating ENSE00002796125 exon, *PLIN3B*) were extracted by regtools73 and a ratio Junction3/(Junction3+Junction4) was calculated for each tissue.

### Sample preparation and proteomic analysis of AON treated cells for PLIN3B detection

Washed cell pellets of AON treated cells (see **morpholino antisense oligonucleotide treatment**) were lysed by resuspension in 120 μl of 4% SDS, 20 mM DTT, 100 mM Tris, pH 7.5) and treatment at 75°C for 10’. After sonication, cleared extracts were loaded on a 12 % mini polyacrylamide gel, migrated about 5 cm and stained by Coomassie blue. Proteins migrating between 35-55 kDa were excised and digested with sequencing-grade trypsin as described (Shevchenko et al., 2007). Extracted tryptic peptides were dried and resuspended in 0.1% formic acid for MS analysis.

MS analysis was performed by data dependent acquisition on a Fusion Tribrid Orbitrap mass spectrometer (Thermo Fisher Scientific). LC conditions were as described above. Full MS survey scans were performed at 120’000 resolution. A data-dependent acquisition method controlled by Xcalibur software (Thermo Fisher Scientific) was used that optimized the number of precursors selected (“top speed”) of charge 2^+^ to 5^+^ while maintaining a fixed scan cycle of 0.6s. Peptides were fragmented by higher energy collision dissociation (HCD) with a normalized energy of 32%. The precursor isolation window used was 1.6 Th, and the MS2 scans were done in the ion trap. The *m/z* of fragmented precursors was then dynamically excluded from selection during 60s. After a standard DDA analysis, samples were also run with a customized method targeting the peptide of interest (QEQSYFMETVK, m/z=695.32 (2^+^)) spanning the PLIN3 splice site, predicted to exist only in the PLIN3B isoform. The detectability, m/z and retention time of the peptide was later confirmed by analysing a synthetic peptide with the same MS method.

MS data were analysed using Mascot 2.8 (Matrix Science, London, UK) set up to search the human database as described above, a custom database including contaminants as well as the PLIN3B sequence. Trypsin (cleavage at K, R) was used as the enzyme definition, allowing 2 missed cleavages. Mascot was searched with a parent ion tolerance of 10 ppm and a fragment ion mass tolerance of 0.02 Da. Carbamidomethylation of cysteine was specified in Mascot as a fixed modification. Protein N-terminal acetylation and methionine oxidation were specified as variable modifications. LC-MS traces were manually inspected to ascertain signal for the targeted peptide.

### Ribosome sequencing database analysis

Ribosome sequencing data was obtained from Trips-Vitz at https://trips.ucc.ie/ (Kiniry et al., 2021) (Gencode_v25), querying ENST00000589163.5 (*K7ERZ3*) and ENST00000221957.9 (*PLIN3A*) ribosome coverage data from the 654 data files available in the database.

### Protein structure prediction and visualization

PLIN3A-ALFA, PLIN3B and PLIN3B-ALFA protein structure was predicted with AlphaFold2 (Herráez, 2006). Protein structures were visualized with UCSF ChimeraX (Meng et al., 2023). Predicted aligned error of the protein prediction was visualized with PAE viewer (Elfmann and Stulke, 2023).

### Transmission Electron microscopy

Cells grown on Mattek gridded dishes dish were fixed in 2.5% glutaraldehyde (EMS, Hatfield, PA, US) in Phosphate Buffer (PB 0.1M pH 7.4) (Sigma) during 1 h at RT. Then they were directly postfixed by a fresh mixture of osmium tetroxide 1% (EMS, Hatfield, PA, US) with 1.5% of potassium ferrocyanide (Sigma, St Louis, MO, US) in PB buffer during 1 h at RT. The samples were washed three times in distilled water and dehydrated in ethanol solution (Sigma, St Louis, MO, US) at graded concentrations (30%-10’; 70%-10’; 100%-3퀇10’). This was followed by infiltration in Epon (Sigma) ON, then replaced the day after with fresh resin on the coverslip and finally polymerized for 48 h at 60°C in oven. The coverslips were removed with razor blades, and the flat resin blocks were imaged with a digital microscope (Keyence, Osaka, Japan) to localize cells of interest. The flat block region was cut and glued on an epoxy resin block and ultrathin sections of 50nm were cut on a Leica Ultracut (Leica Mikrosysteme GmbH) and picked up on a copper slot grid 2퀇1mm (EMS, Hatfield, PA, US) coated with a PEI film (Sigma). Sections were poststained with uranyl acetate (Sigma) 2% in water for 10’, rinsed several times with water followed by Reynolds lead citrate in water (Sigma) during 10’ and rinsed several times with water. For the analysis, montages were collected with a pixel size of 4.402nm with a transmission electron microscope Philips CM100 (Thermo Fisher Scientific) at an acceleration voltage of 80kV with a TVIPS TemCam-F416 digital camera (TVIPS GmbH, Gauting, Germany). Montage alignments were performed using Blendmont command-line program from the IMOD software (Kremer et al., 1996).

### FluorNanogold-CLEM

Cell grown on Mattek gridded dishes (Mattek 35 mm Dish | No. 1.5 Gridded Coverslip | 14 mm Glass Diameter) were fixed in 4% paraformaldehyde and 2.5% glutaraldehyde (EMS, Hatfield, PA, US) in Phosphate Buffer (PB 0.1M pH 7.4) (Sigma, St Louis, MO, US) during 30’ at 37°C. After rinsing in PB, residual aldehydes were reduced with 1mg/mL Sodium borohydride (Sigma-Aldrich, 452882) in PB for 10’. Permeabilization was performed based on (Knott et al., 2009). In brief, cells were incubated in cryoprotecion solution (20% DMSO, 2% glycerol in PB) and plunged in liquid nitrogen, then thawed by addition of fresh cryoprotection solution. Freeze-thaw cycle was performed thrice. Cells were blocked in 5% FBS, 2% BSA in PB for 30’, then incubated with first antibodies for 1 h, washed thrice and incubated with secondary antibodies diluted in blocking solution for 1 h. Cells were washed thrice, fixed 5’ in 4% formaldeyhe-PB, then stored in PB. Confocal imaging was performed with Stellaris 8 CLSM at 630X magnification. 3D reconstruction was performed with Imaris (Oxford Instruments, High Wycombe, UK, RRID:SCR_007370). Silver enhancement was performed with Aurion R-Gent Silver Enhancement EM Kit (Aurion, Wageningen, Netherlands, 500.033) for 5’ according to manufacturer’s protocol. After several washes, samples were postfixed by a fresh mixture of osmium tetroxide 0.5% (EMS) in PB buffer during 15’ at RT. The samples were then washed thrice in distilled water and dehydrated in graded series of ethanol solutions (Sigma) (30%-10’; 70%-10’; 100%-3퀇10’). This was followed by infiltration in Epon (Sigma), with a mixture of ethanol and Epon at 1:1 for 30’, Epon 100% for 30’, Epon 100% ON, then replaced the day after with fresh resin on the coverslip and finally polymerized for 48 h at 60°C in the oven. The coverslips were manually removed with a razor blade, and the flat resin blocks were imaged with a digital microscope (Keyence) to localize cells of interest and visualise the grid pattern at the surface of the resin. The region of interest was cut and glued on a flattened epoxy resin block and 100 nm-thick serial sections were cut on a Leica EM UC7 Ultramicrotome (Leica Mikrosysteme GmbH) and picked up on a copper slot grid 2퀇1mm (EMS) coated with a PEI film (Sigma).

Micrographs were acquired using a Talos L120C transmission electron microscope (Thermo Fisher Scientific) at an acceleration voltage of 120kV, equipped with a CETA camera (Thermo Fisher Scientific), at magnifications ranging from 8’500-36’000x, and pixel size up to 402.9 pm. Another part of the micrographs were acquired using a Tecnai G2 Spirit transmission electron microscope (Thermo Fisher Scientific) at an acceleration voltage of 80kV, equipped with a TVIPS camera (Tietz Video and Image Processing Systems), at magnifications ranging from 1250-30’000X, and a pixel size up to 0.7nm.Serial images alignment as well as 3D correlation with the confocal stacks was done using ImageJ2 (Rueden et al., 2017).

### Flow Cytometry

Trypsinized cell pellets were resuspended in ice-cold PBS supplemented with 2% v/v FBS, washed one time in PBS-FBS and analysed with Cytoflex S (Beckman coulter, Brea, CA, USA). FCS files were analysed with Floreada.io (https://floreada.io). In brief, fluorescence bleed through compensation was performed using the Floreada.io autospill function, followed by living cell population selection with forward scatter (FSC) vs side scatter (SSC). A subsequent FSC area vs FSC height gating was used to select single cells. Transfected cell populations were gated by excluding all points having BFP fluorescence intensity values comparable to the non-transfected control samples. Median Fluorescence intensity (MFI) for every channel fluorescence area was then measured and values analysed.

### Statistical analysis

All statistical analyses were performed using GraphPad Prism 9 (GraphPad Software, La Jolla, USA). Data are presented as mean ± standard error of the mean (SEM) unless otherwise specified. In truncated violin plots, lines represent median and quartiles. All reported sample sizes represent biological replicates unless otherwise stated.

Normality of data was assessed using the Shapiro–Wilk test, and equality of variances was evaluated using Brown–Forsythe test. For more than two group comparisons, if normality assumption was met, One-way ANOVA was performed with Tukey HSD multiple comparison test. If normality assumption was not met, Kruskal-Wallis test was performed with Dunn’s multiple comparison test. For comparisons between two independent groups, two-tailed unpaired Student’s *t*-tests were used when assumptions of normality and equal variance were met. If the assumption of normality was not met, Mann–Whitney U test was used. Multispectral analyses between the 3 conditions (Ctrl, PLIN3A, PLIN3B) were compared with mixed-effects analysis, with the Geisser-Greenhouse correction. To correct for multiple comparisons by controlling the False Discovery rate two-stage linear step-up procedure of Benjamini, Krieger and Yekutieli were performed with individual variances computed for each comparison. For all tests, a *P* value of < 0.05 was considered statistically significant.

